# SkeletAge: Transcriptomics-based Aging Clock Identifies 26 New Targets in Skeletal Muscle Aging

**DOI:** 10.1101/2025.07.28.667277

**Authors:** Muhammad Ali, Fei Li, Manpreet Katari

**Affiliations:** Department of Biology, New York University, New York, NY, 10003, United States

**Keywords:** Ageprint, Biological Aging, Epigenetic Clock, Longevity Drug Discovery, Skeletal Muscle

## Abstract

Identifying the set of genes that regulate baseline healthy aging – aging that is not confounded by illness – is critical to understating aging biology. Machine learning-based age-estimators (such as epigenetic clocks) offer a robust method for capturing biomarkers that strongly correlate with age. In principle, we can use these estimators to find novel targets for aging research, which can then be used for developing drugs that can extend the healthspan. However, methylation-based clocks do not provide direct mechanistic insight into aging, limiting their utility for drug discovery. Here, we describe a method for building tissue-specific bulk RNA-seq-based age-estimators that can be used to identify the *ageprint*. The ageprint is a set of genes that drive baseline healthy aging in a tissue-specific, developmentally-linked fashion. Using our age estimator, SkeletAge, we narrowed down the ageprint of human skeletal muscles to 128 genes, of which 26 genes have never been studied in the context of aging or aging-associated phenotypes. The ageprint of skeletal muscles can be linked to known phenotypes of skeletal muscle aging and development, which further supports our hypothesis that the ageprint genes drive (healthy) aging along the growth-development-aging axis, which is separate from (biological) aging that takes place due to illness or stochastic damage. Lastly, we show that using our method, we can find druggable targets for aging research and use the ageprint to accurately assess the effect of therapeutic interventions, which can further accelerate the discovery of longevity-enhancing drugs.

## Introduction

Historically, aging has been largely accepted as a part of life. However, recent advances in geroscience have helped usher in a social paradigm shift towards accepting aging as more of a disease (de Magalhães, 2014) that, at the very least, warrants investigation into its molecular architecture and, at most, needs a cure.

Over the years, researchers have uncovered lifestyle interventions that may extend lifespan, such as caloric restriction (CR) and exercise. However, CR, which shows optimistic results in model organisms, does not translate very efficiently in humans, with an average lifespan extension of up to 5%, contingent on the age at which an individual starts CR (Flanagan et al., 2020). There is some evidence that CR may reduce the pace of aging by about 2 – 3% (Waziry et al., 2023), which could offer promising dividends when compounded over the years. However, given the low returns, it is still far from being a readily adaptable strategy. On the other hand, regular moderate exercise has shown numerous benefits ranging from neuroprotection (Allen et al., 2021; Bettio et al., 2020; Stillman et al., 2020) to reduction in inflammation (Lavin et al., 2020; Radak et al., 2008), protection from metabolic (Thyfault & Bergouignan, 2020) and cardiovascular disorders (G. Li et al., 2020) and more. However, retrospective analyses have shown that regular exercise can only extend lifespan by up to 4 years (Reimers et al., 2012); spending 300 – 600 minutes every week on voluntary moderate exercise for over 30 years only reduces the risk of all-cause mortality by up to 31% (Lee et al., 2022).

Given the meager effect of lifestyle interventions on aging, it is no surprise that even with an ideal “anti-aging” lifestyle, we would still expect an individual to age. This gives rise to the hypothesis that aging is not just the accumulation of stochastic damage over the years but is also developmentally-coded into our DNA (de Magalhães, 2012). Thus, it stands to reason that, like any other biological process, aging – at least to some degree – must also be under the control of certain genes. If we could identify these genes, we can not only begin to understand the mechanisms of aging but also start developing pharmacological interventions against aging that could potentially extend the lifespan beyond the current best lifestyle-based interventions’ capacity. Studies in model organisms, such as *C. elegans* (Kenyon et al., 1993; Uno & Nishida, 2016) and mice (Dubnov et al., 2024; Prochownik & Wang, 2023; Ruetz et al., 2024), have demonstrated that genes can greatly influence the lifespan of an organism. Thus, it is reasonable to presume that an underlying developmental-genetic-footprint regulates how we age. We hypothesize that this *ageprint* is a tissue-specific unchanging set of co-expressed (and perhaps co-regulated) genes whose expression is the main driver of “healthy” baseline aging – the age-associated phenotypes that appear despite living a healthy lifestyle and not having any (chronic) illness. One of the main reasons we believe the *ageprint* is tissue-specific is the fact that different tissues do not always express the same genes at the same levels. Moreover, there is evidence that tissues do not age at the same rate (Oh et al., 2023; Slieker et al., 2018; Tian et al., 2023; Tuttle et al., 2020), which further suggests that the underlying mechanisms of aging differ from tissue-to-tissue. Thus, identifying a tissue’s ageprint is pivotal to understanding and eventually preventing (biological) aging in said tissue. Ideally, the best way to identify the ageprint would be doing thousands, if not hundreds of thousands, of *in vivo* genetic screens. However, there are thousands of genes in a single organism, and a multitude of them likely regulates aging. Therefore, in a race against time, using knock-out screens and then translating them into aging therapeutics may take decades, if not centuries. Several groups have tried to look at differentially expressed genes (DEGs) between younger and older people to identify the mechanisms of aging (Avelar et al., 2020; B. Ko & Van Raamsdonk, 2023; Lopes et al., 2022; Pan et al., 2020; Peters et al., 2015). However, we believe that the ageprint may not be differentially expressed between older and younger people. Differential expression represents drastic changes in gene expression, which are more likely to be associated with the damage-accumulating model of aging, rather than the developmental model of aging. Thus, we believe that the ageprint will likely demonstrate relatively stable expression over the ages because it is driving aging that cannot be accounted for by disease, damage, or perturbations; it is the innate aging that cells are programmed to go through. Thus, one of the fastest ways to identify the ageprint is to leverage machine learning – specifically “aging clocks” – and find genes that explain aging in healthy people at both older and younger ages. Aging clocks – or age-estimators – are essentially machine learning models that can predict chronological (or biological) age based on a set of biomarkers, such as telomere lengths (Blackburn et al., 2006), DNA methylation (Hannum et al., 2013; Horvath, 2013), transcriptional profiles (Holly et al., 2013; Peters et al., 2015; Ren & Kuan, 2020), IgG glycosylation level (Krištić et al., 2014), quantitative proteomic data (Menni et al., 2015), metabolite levels (Hertel et al., 2016; Menni et al., 2013). Some clocks combine several biomarkers to create composite *biological aging* scores (Belsky et al., 2015; Levine et al., 2018; Sebastiani et al., 2017) and can even predict all-cause mortality (Lu et al., 2019). Fascinating as it is, these clocks can also be used to identify the most important biomarkers for aging from a huge set of biomarkers – such as all the genes in a genome – using feature selection methods. Thus, it is reasonable to presume that any set of biomarkers that strongly correlates with aging and can explain aging both at younger and older ages may offer insight into the mechanisms of (healthy) aging.

Currently, the gold standard for age-estimators is methylation-based clocks (Jylhävä et al., 2017), commonly known as epigenetic clocks, DNAm clocks, or epigenetic age estimators. These clocks tend to have low errors and usually a high correlation between the predicted epigenetic age and chronological age. Epigenetic clocks were first popularized by the Hannum clock (Hannum et al., 2013) and Horvath’s clock (Horvath, 2013). Since then, many different clocks have been developed, with PhenoAge (Levine et al., 2018) and GrimAge (Lu et al., 2019) being two of the most popular second-generation clocks that can predict biological age and all-cause mortality, respectively. These clocks are based off one common principle: methylation levels at certain CpGs (herein referred to as “clock CpGs”) across the genome are linearly correlated with age (Bocklandt et al., 2011). This phenomenon allows us to build robust (epigenetic) age estimators using methylation data to precisely predict an individual’s chronological (or biological) age and measure potential epigenetic age acceleration or deceleration (Horvath & Raj, 2018). Moreover, these clocks can also help us identify which CpGs are associated with aging. However, the biggest problem with epigenetic clocks is the lack of direct mechanistic relevance of CpGs (Bell et al., 2019) in aging; we do not know how CpG methylation directly translates to processes that influence aging. Different groups have tried to gain a mechanistic understanding of the importance of clock CpGs by trying to find associations with genes (Koch & Wagner, 2011) or genomic locations (A. Li et al., 2022; Yan et al., 2020), or they tried mapping the location of these CpGs to promoters and enhancers (Hannum et al., 2013; Horvath, 2013) to gain a better biological insight into the functional relevance of these clock CpGs in aging. However, these correlations only serve as a surrogate for a real mechanistic explanation.

On the other hand, biological clocks that use biomarkers whose mechanism can be more easily interpreted, such as clocks based on transcriptomics (Holly et al., 2013; Peters et al., 2015; Ren & Kuan, 2020), proteomics (Menni et al., 2015), metabolomics (Hertel et al., 2016; Menni et al., 2013), or even the gut microbiome (Gopu et al., 2024), are usually not as precise as DNAm clocks (A. Li et al., 2022). That could be because people try to build pan-tissue clocks. Given that every tissue’s underlying ageprint may differ, it is easy to see why most of these clocks may not achieve great precision when trained using many different types of tissues – the number of non-CpG biomarkers that can explain aging in all tissues might be very small. The tissue-specific noise in the data may add a lot of variance that throws off the model. Thus, to readily identify the ageprint of a tissue, we need to move away from pan-tissue clocks towards robust tissue-specific clocks that are built using easily-interpretable biomarkers. This would allow us to derive mechanistic insights into a particular tissue’s aging directly from the clocks and help us identify new targets for drug development.

To explore this hypothesis, we built a transcriptomics-based clock for skeletal muscles: *SkeletAge*. We chose skeletal muscles because we wanted to capture the genes driving true baseline aging – the *ageprint* – and muscles exhibit primary sarcopenia, an age-associated decline in muscle mass that any other disease or disorder cannot explain (Santilli et al., 2014). This is an example of “healthy” aging, i.e., when a person who is free of any major diseases or conditions and follows a healthy lifestyle experiences age-related phenotypes. The existing transcriptomics-based age estimators either did not explore tissue-specificity and focused more on generalizability (Peters et al., 2015) or were trained on post-mortem samples’ RNA-seq data (Ren & Kuan, 2020), which may not accurately reflect the baseline transcriptomic profile of the antemortem tissue (Franz et al., 2005; Schuurs et al., 2006; Yeh et al., 1999). To the best of our knowledge, there are no transcriptomics-based age estimators specifically for skeletal muscles built using RNA-seq data exclusively from healthy people. Thus, we trained SkeletAge on skeletal muscle RNA-seq data specifically from healthy people over a broad age range (19 – 90 years). We hypothesized that due to the nature of SkeletAge’s training set, we would be able to identify the genes that form the ageprint of skeletal muscles because these genes would explain healthy aging at both younger and older ages, which suggests that these genes *imprint* aging into the skeletal muscle tissue.

In line with our hypothesis, SkeletAge performed remarkably and showed a strong correlation between the predicted and chronological age (0.91). It also narrowed down the skeletal muscles’ ageprint to 128 genes that seem to be central to aging skeletal muscles; 26 of these genes have never been studied in the context of aging or age-associated phenotypes before. The ageprint identified by SkeletAge can be linked to both development and known skeletal muscle aging phenotypes. Additionally, we found that 23 out of the 128 genes were present in the druggable genome. Of the drugs associated with the druggable genes, 4 approved drugs had already shown promising results in age-related studies. Hence, we demonstrate that tissue-specific RNA-seq-based age-estimators can not only pinpoint a tissue’s ageprint but also discover novel genetic targets for aging. This opens doors to a powerful new way of using tissue-specific age estimators to identify a tissue’s ageprint, which will help in not only untangling the web of genetic mechanisms that influence tissue-specific aging but also accelerate the development and testing of aging therapies by identifying druggable targets and providing a quick, intelligible readout in the form of direct transcriptomic changes, respectively.

## Results

### The right training set for *ageprint* detection

To identify the genes of interest in skeletal muscle aging, we built a transcriptomics-based skeletal muscle aging clock called *SkeletAge*. We wanted to capture the baseline tissue-specific signature of aging – the *ageprint*. Given that we hypothesized the ageprint to be a genetic signature that explains healthy aging at both older and younger ages and is selected based on a machine learning model, it was imperative that we chose the right training set. If the people in our training set were not healthy (or too healthy – such as professional athletes), they could have confounded the feature selection process of the model. Therefore, by analyzing multiple studies on healthy aging genomics (Robinson et al., 2017; Tumasian et al., 2021), we devised a set of criteria for exclusion from the training set: BMI >20 and <30 kg/m², no regular exercise (more than 20 minutes twice a week), or history of professional athleticism or training. The participants in the training set should also be free of any major chronic illnesses or conditions such as heart disease, lung problems, metabolic disorders, neurological pathologies, or cancer. They must not have any physical or cognitive impairment and must not have undergone a serious medical procedure such as surgery. Additionally, they must not be on any medication needed to manage chronic conditions. The individuals need to be alive at the time of sample collection because of the marked differences between the antemortem transcriptome and the thanatotranscriptome of skeletal muscles (Sanoudou et al., 2004). Based on this selection criteria, we trained an elastic net model (λ = 19.15027 | α = 0.1) using transcriptomics data of healthy subjects from three different studies that closely met this criterion: GSE164471, GSE97084, and GSE144304. In total, we had 236 samples that we used to train our clock. The individuals comprised a broad age range (19 – 90 years old | mean age: 52 years | SD: ±24.6 years) and were reported to be healthy when their samples were taken (Figure 1).

**Figure 1:**
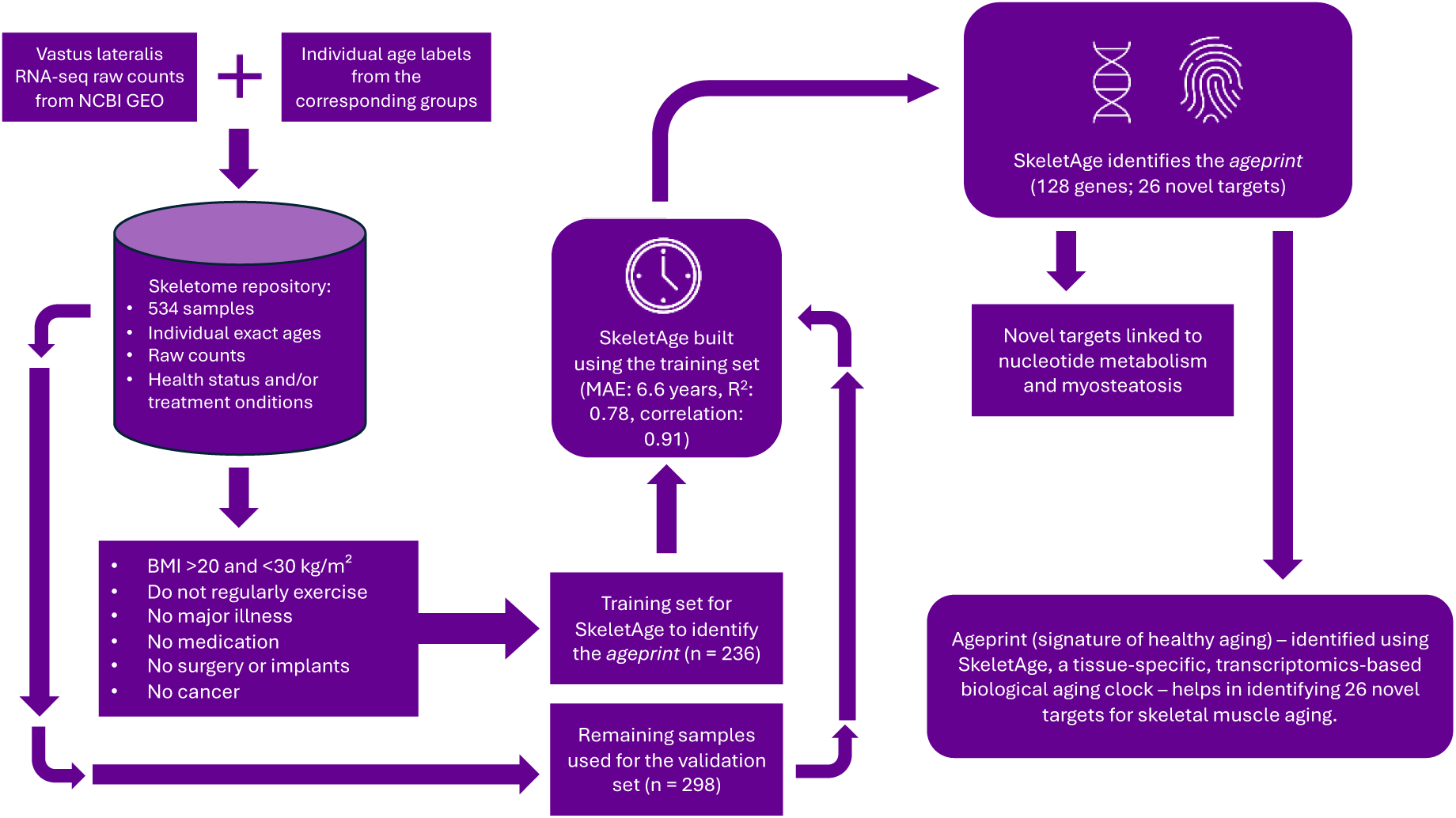
Creating SkeletAge and identifying the ageprint. We created SkeletAge by taking RNA-seq data of vastus lateralis samples on NCBI GEO and training an elastic net model to predict ages using this gene expression data. This allowed us to build a tissue-specific clock that can identify the ageprint. Using the ageprint, we identified 26 novel genetic targets for skeletal muscle aging that have never been reported in the context of aging research. Creating and testing SkeletAge took about 534 samples, for which accurate age labels had to be acquired from the corresponding authors. Therefore, to accelerate future aging research, we have compiled our dataset, including the metadata with accurate age labels and the raw counts for each sample (along with its GEO sample accession), into a repository called *SkeletAge*, which is accessible through this link: https://github.com/mali8308/SkeletAge/.

### Performance metrics of SkeletAge

Next, we sought to test how well SkeletAge could predict someone’s chronological age. SkeletAge performed decently enough in the training set and strongly correlated with chronological age (MAE: 6.55 years | R^2^: 0.90 | Correlation: 0.96). We further validated SkeletAge using another 298 samples from 8 different studies (GSE196387, GSE20543, GSE157988, GSE151066, GSE58608, GSE159217, GSE200398, GSE186045). The samples in the validation set exhibited various ages, conditions, and/or treatments. For instance, there were athletes, people who were obese, or people suffering from end-stage osteoarthritis. We chose a dataset with a wide variety of ages and conditions to see whether our clock can predict the chronological age precisely, regardless of health status. Being able to do so would suggest that it predicts age based on the genes that drive baseline aging and are likely involved in the developmentally programmed aging mechanisms that are non-redundant with the stochastic damage-associated pathways of aging. Even for this widely diverse dataset, SkeletAge performed well and demonstrated a strong correlation with the chronological age (MAE: 6.62 | R^2^: 0.79 | Correlation: 0.91). Using a Wilcoxon signed rank test, we found that the difference between the samples’ predicted and real ages was not statistically significant (V = 21723 | p-value = 0.7108). This suggests that SkeletAge is indeed capturing the ageprint of skeletal muscles that drives baseline aging regardless of the individual’s health status. The scatter plot for the validation set’s predicted ages and the best fit line for the validation set can be found in Figure 2A. Detailed information about all the conditions and groups used for the training and the testing is present in Table S1.

**Figure 2:**
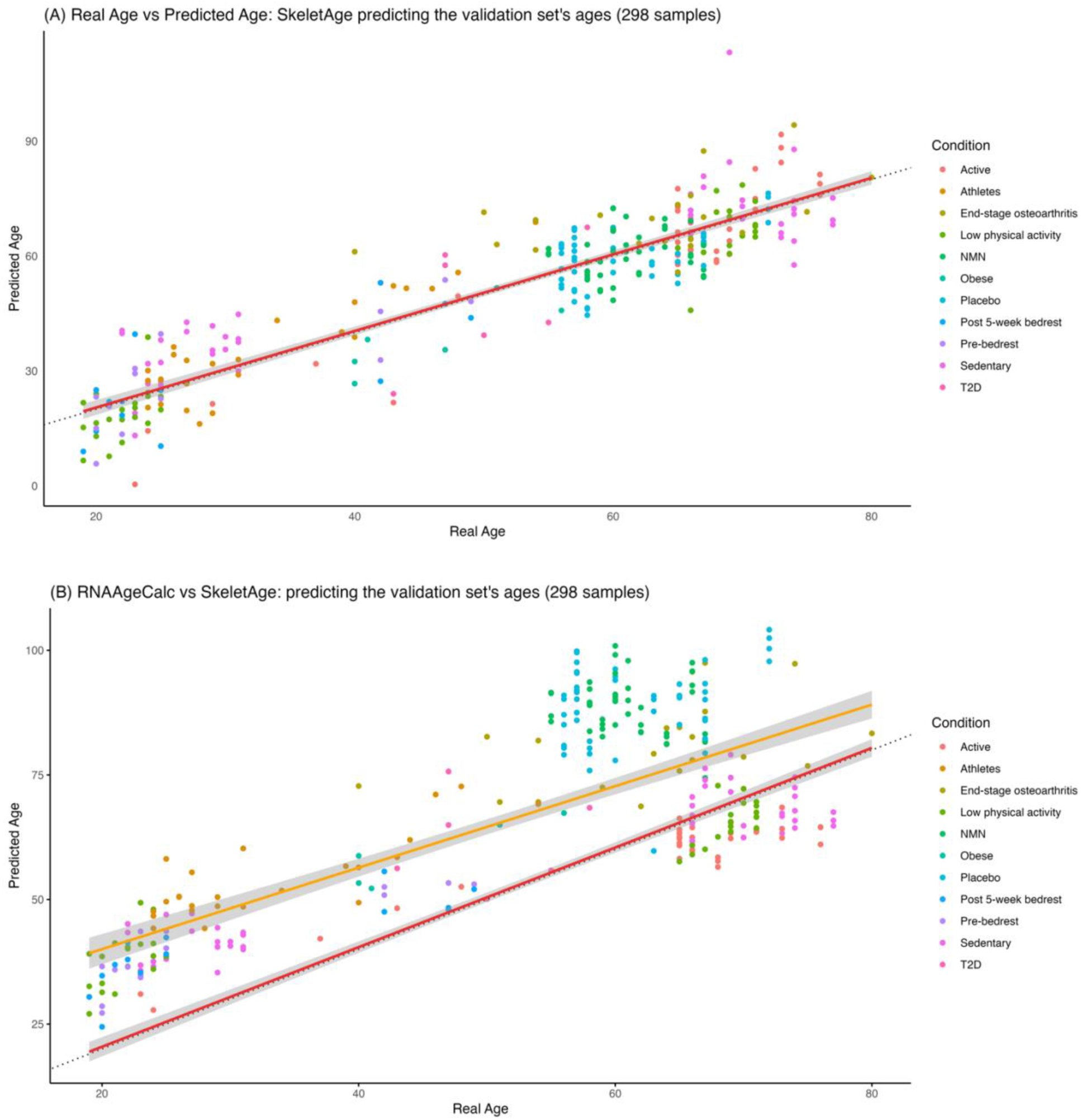
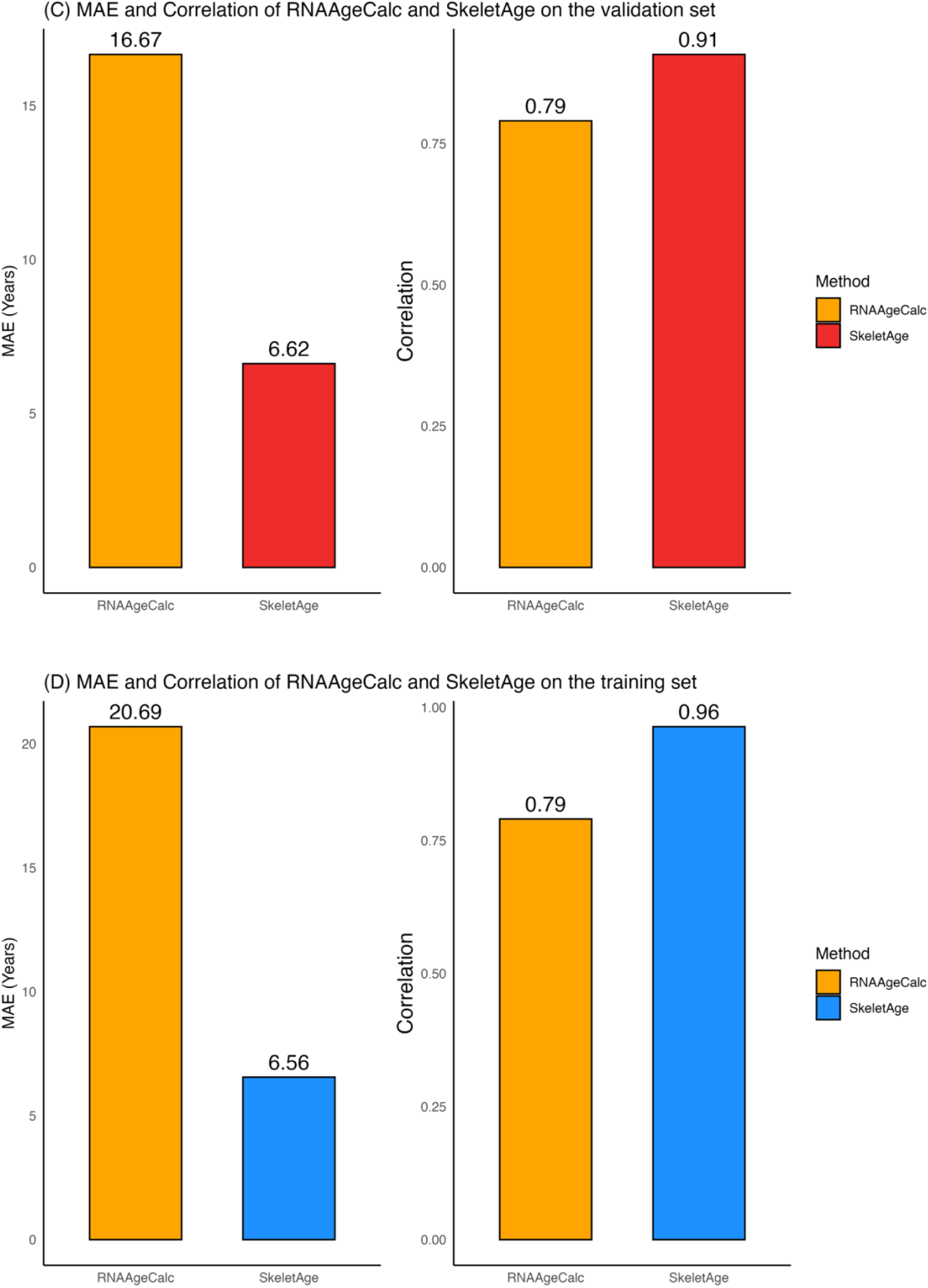
Comparison between SkeletAge and RNAAgeCalc. (A) shows the predicated ages measured by SkeletAge, whereas (B) shows the predicted ages (RNAAge) as measured by RNAAgeCalc. Each point on the scatterplot is a sample from the validation set (n = 298). The color of the points provides the health status or treatment groups. The red line is the line of best fit generated by SkeletAge, and the orange line is the line of best fit generated by RNAAgeCalc. Both models are predicting the ages for the validation set. The dotted line represents a 1:1 correlation between the chronological and the predicted ages. (C) shows the comparison of the mean absolute error (MAE) and Pearson correlation between RNAAgeCalc and SkeletAge when they are used on the validation set (n = 298). (D) is a comparison of the mean absolute error (MAE) and Pearson correlation between RNAAgeCalc and SkeletAge when they are used on the training set (n = 236).

### Comparison of SkeletAge with RNAAgeCalc in skeletal muscles

SkeletAge is a tissue-specific age-estimator trained to predict the ageprint – the baseline signature of healthy aging – in order to identify targets for tissue-specific aging therapies. To compare whether training our model using data exclusively from healthy people offered a significant advantage over a pan-tissue RNA-based age estimator trained on data with less stringent constraints for their exclusion criteria, we compared SkeletAge with RNAAgeCalc (Ren & Kuan, 2020). We installed the RNAAgeCalc package for R using Bioconductor and found that for our validation set, RNAAgeCalc performed worse (MAE: 16.7 years | R^2^: −0.16 | Correlation: 0.75; 95% CI for Correlation: 0.69 – 0.79), compared to SkeletAge (MAE: 6.62 | R^2^: 0.79 | Correlation: 0.91; 95% CI for Correlation: 0.89 – 0.93), as shown in Figure 2B and 2C. Since our validation set was very diverse with many conditions and/or treatments, we assumed that the phenotypic variation may be confounding RNAAgeCalc’s predictions. Therefore, we decided to test RNAAgeCalc on SkeletAge’s training set, which consisted of individuals reported to be healthy (Figure 2D). Here, RNAAgeCalc performed relatively better (MAE: 20.7 years | R^2^: 0.58 | Correlation: 0.79; 95% CI for Correlation: 0.74 – 0.83) but was unsurprisingly outmatched by SkeletAge (MAE: 6.56 | R^2^: 0.90 | Correlation: 0.96; 95% CI for Correlation: 0.95 – 0.97). This suggests that RNAAgeCalc gets skewed by confounders such as the clinical phenotypes of different individuals and thus does not use the ageprint for age prediction as SkeletAge does. This is likely due to the fundamental differences in the training sets of the two models. While we trained SkeletAge using data from people who were healthy and alive at the time of sample collection, RNAAgeCalc was built using post-mortem bulk RNA-seq data from GTEx (Carithers et al., 2015; Lonsdale et al., 2013). Due to the post-mortem nature of tissue collection, gene expression levels may not accurately reflect the true phenotypic nature of the antemortem tissue (Franz et al., 2005; Schuurs et al., 2006; Yeh et al., 1999). The skeletal muscle thanatotranscriptome shows marked differences from the antemortem gene expression patterns (Sanoudou et al., 2004), which may explain why RNAAgeCalc cannot identify the ageprint of a tissue. We used Wilcoxon signed rank tests to validate the differences between RNAAgeCalc and SkeletAge. RNAAgeCalc’s predicted age – RNAAge – was significantly different from the chronological ages of the samples in both the training set (V = 9253 | p-value = 6.649e^−06^) and the validation set (V = 3494 | p-value = 1.742e^−36^). On the other hand, the difference between the ages predicted by SkeletAge and the chronological ages of the samples was not significant in both the training set (V = 13902 | p-value = 0.9389) and the validation set (V = 21723 | p-value = 0.7108). Additionally, we found that the difference in the ages predicted by SkeletAge and RNAAge was also significant for both the training (V = 20466 | p-value = 6.646e^−10^) and the validation sets (V = 39880 | p-value = 2.913e^−32^). SkeletAge’s validation set error was also smaller than the error reported for the peripheral blood-based transcriptomic age-predictor built by Peters et al.: 6.6 years for SkeletAge compared to 7.8 years for the age-estimator built using peripheral blood data (Peters et al., 2015). This further highlights the utility of moving from generalizable blood-based clocks to more precise tissue-specific age-estimators that can capture the ageprint of a specific tissue, which can then be used to understand the mechanisms of aging and discover potential targets for aging therapies.

### SkeletAge predicts 26 novel targets in skeletal muscle aging

Next, we sought to understand what genes SkeletAge was using to calculate the biological age. This would allow us to identify skeletal muscles’ ageprint. To do that, we obtained the genes associated with the 129 non-zero coefficients used by the model. We manually verified that 102 of the 129 genes have previously been reported to be either involved – or studied in the context of – aging or aging-associated phenotypes (Table S3); all the genes and their respective coefficients are present in Table S2. Out of the remaining genes, one gene (EntrezID: 107986257) had its record withdrawn from NCBI because it did not appear in a later iteration of the genome. The remaining 26 genes – to the best of our knowledge – have never been studied in terms of aging, age-associated phenotypes, or skeletal muscle aging; these 26 novel targets are present in Table 1. We further confirmed the centrality of these genes in capturing the ageprint in skeletal muscle tissue by running a principal component analysis (PCA) using the normalized counts of these genes to separate the samples into different clusters. Unsurprisingly, the samples clustered into distinct clusters based on their age groups, regardless of whether we used our training data (Figure 3A; healthy individuals), the validation set (Figure 3C; various conditions), or the entire dataset (Figure 3F; various conditions). An important thing to note here is that the entire dataset contains individuals suffering from conditions such as type-2 diabetes, end-stage osteoarthritis, obesity, and more. Thus, SkeletAge’s genes’ ability to distinctly cluster such a diverse group demonstrates that it robustly captures the signatures of intrinsic aging – the ageprint – which are rooted in the molecular architecture of development that drives aging regardless of health status. Within those 129 genes, 64 were predicted to be decelerating the biological age (β < 0) – referred to as ProLong (promoting longevity) genes from here on – while the remaining 64 (excluding 107986257) – referred to as RedLong (reducing longevity) genes from here on – were predicted to be involved in biological age acceleration (β > 0). Both the ProLong and the RedLong genes’ expression can cluster samples very distinctively in the training set (Figure 3B) and the validation set (Figures 3D and 3E), where the age-based divide is clearly visible. This suggests that these two sets of genes capture the intrinsic aging signatures at a great resolution, even on their own. Interestingly, in the validation set, the PCA for the RedLong genes shows tighter clusters (Figure 3D) compared to the clusters produced by the ProLong genes (Figure 3E). This suggests that the expression pattern of the ProLong genes is more receptive towards phenotypic changes and health status than the RedLong genes because both the ProLong and the RedLong genes produce identical PCA plots for the training set where all samples are healthy (Figure 3B), but in the validation set, where not everyone was healthy, their plots are a little different.

**Figure 3:**
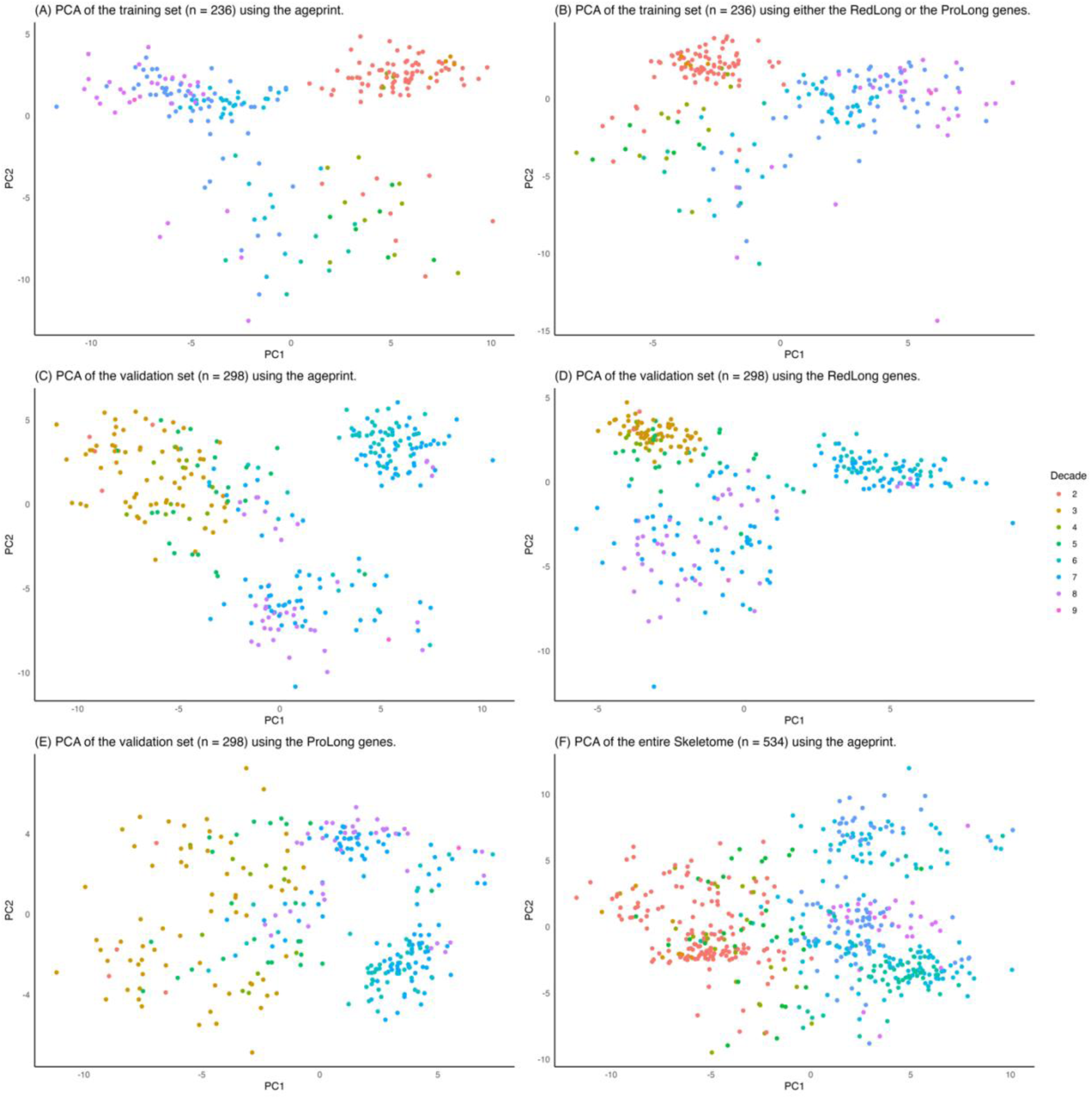
Principal component analysis (PCA) plots generated using the ageprint genes. The plot in (A) shows PCA ran on the training set (only healthy samples). (B) PCA ran on the training set using only the RedLong or the ProLong genes; both produce identical plots, but only one is shown here. (C) The PCA plot generated for the validation set (healthy samples and people suffering from different conditions) uses just the ageprint. (D) The plot generated for the validation set using the RedLong genes, and (E) is the PCA for the validation set using the ProLong genes. (F) shows a PCA plot for the entire Skeletome dataset using the ageprint genes. Each individual point is a sample. The samples are color-coded by the decade they were binned into.

**Table 1:**
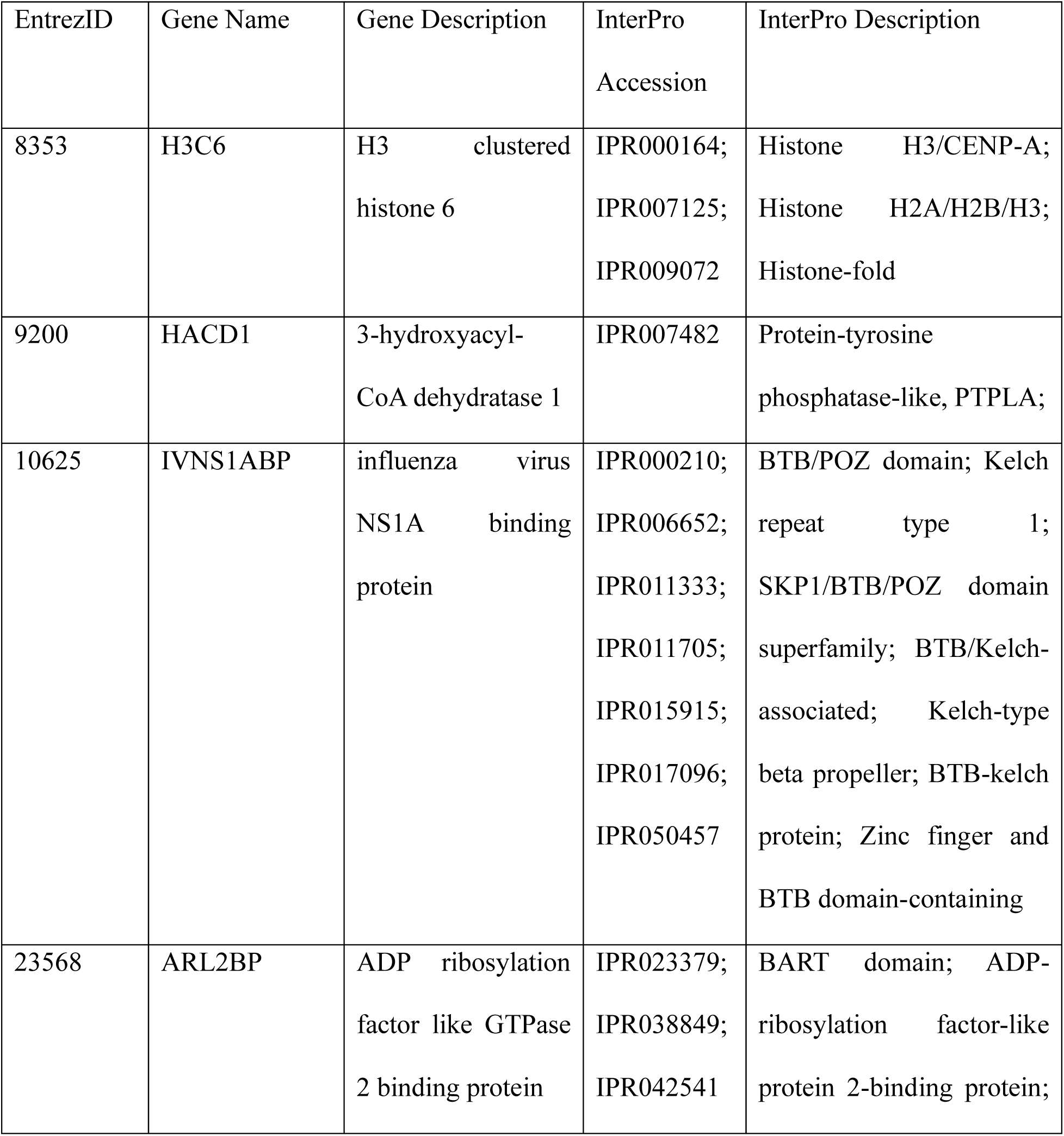
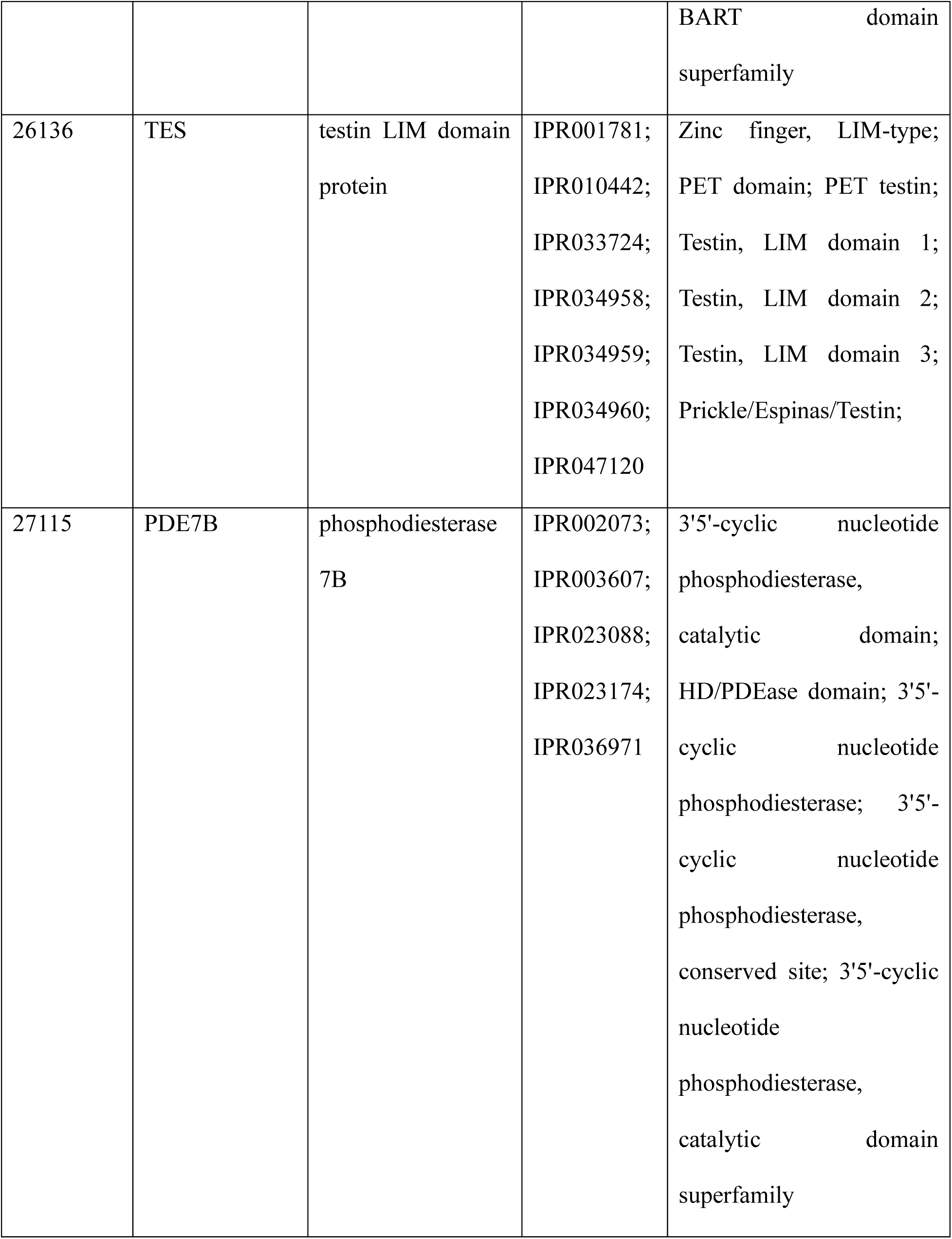
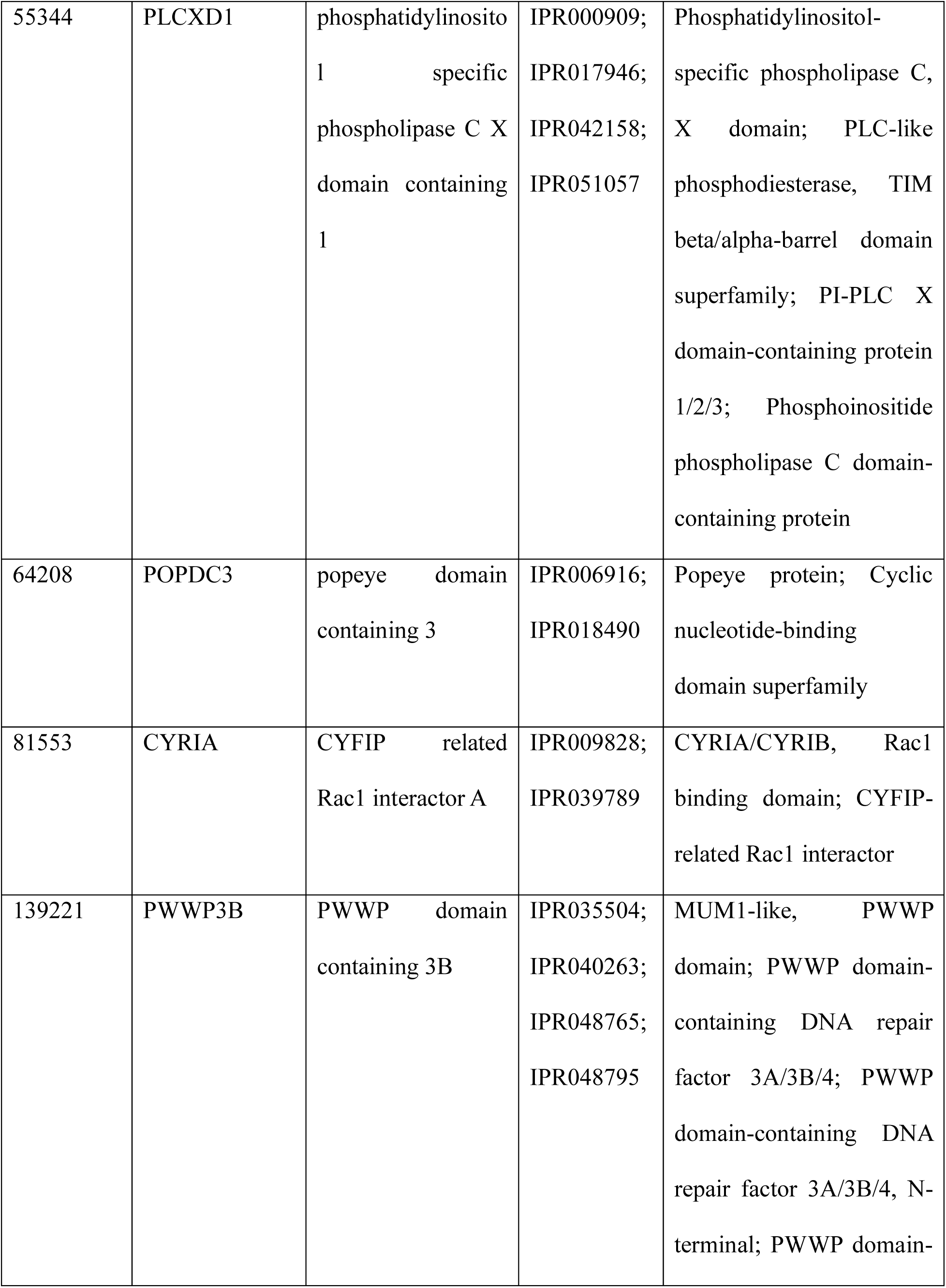
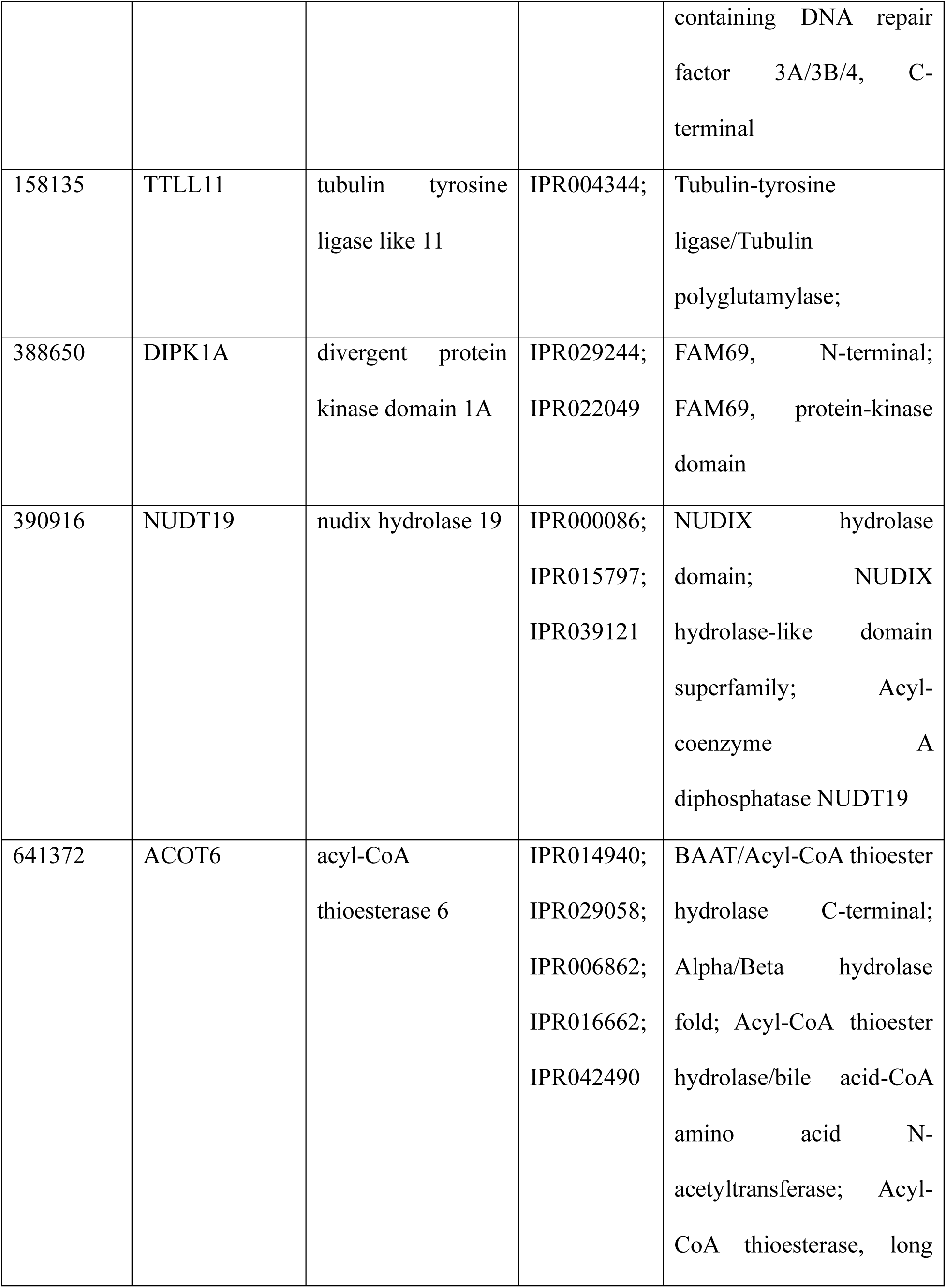
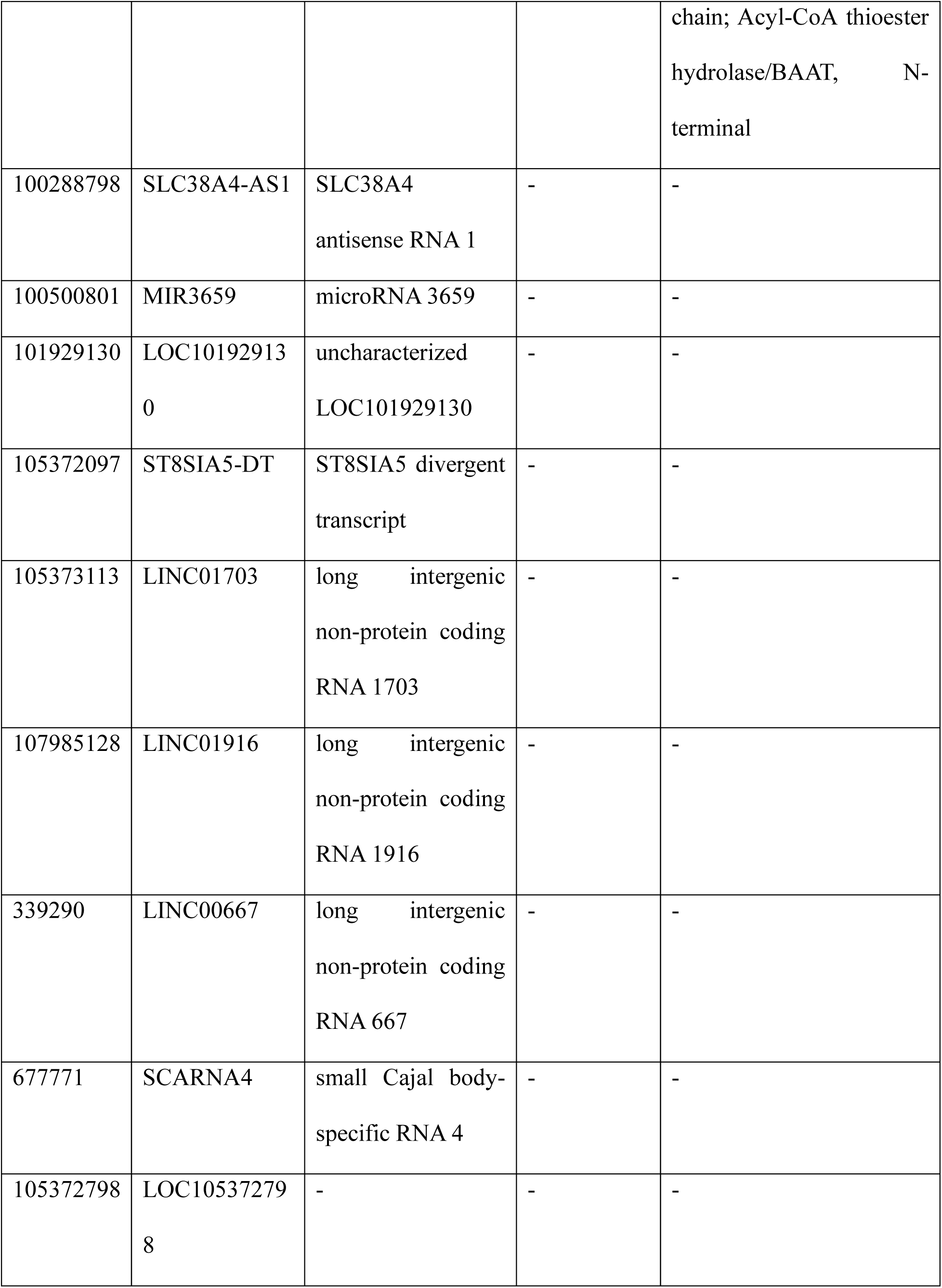
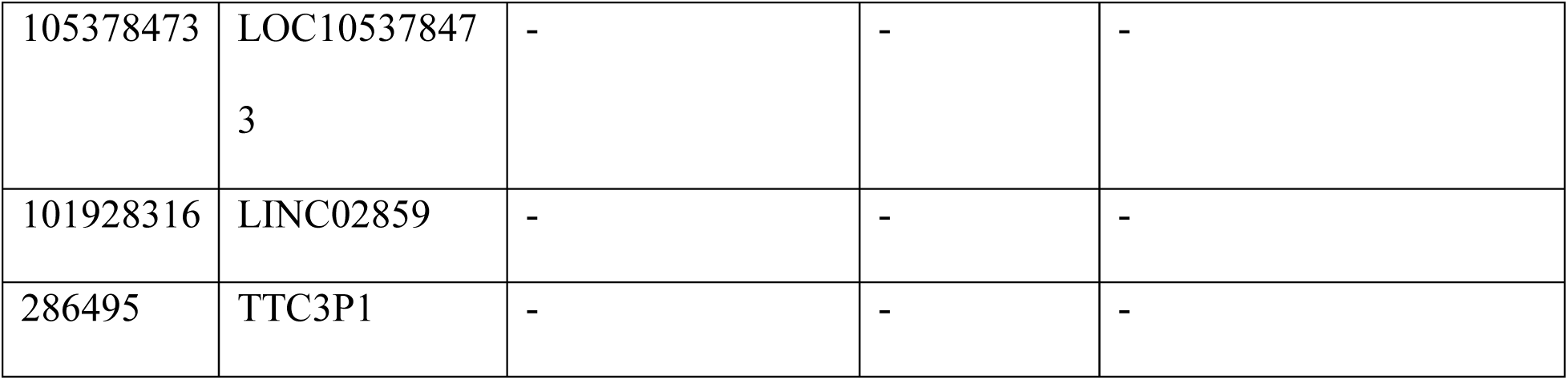
The 26 novel targets predicted by SkeletAge. These 26 genes are a part of the skeletal muscle ageprint (Table S1) identified by SkeletAge. The table contains information about the genes, including InterPro accessions and descriptions for protein-coding genes.

### Relationships between Age, ProLong, and RedLong genes

To further elucidate how the ProLong and RedLong genes were related to each other (and aging), we calculated the correlations between the 128 annotated genes and age. 127 genes were significantly correlated with age (p-value < 0.05), and only *MIR3659* was insignificantly correlated with age (p-value: 0.078). Half of the 128 genes were positively correlated with age, while the other half were negatively correlated with age (Figure 4A). The correlations for either side of the spectrum did not exceed 0.8 or −0.8, respectively. A rather surprising finding here was that all the ProLong genes were negatively correlated with age, while all the RedLong genes were positively correlated with age (Figure 4A). This suggests that genes that accelerate aging undergo an increase in expression as we age. In contrast, the ones that decelerate aging show gradually decreased expression as we age – in principle, that makes sense. To test that theory, we compared the mean log fold changes of ProLong and RedLong genes across the nine different age groups being tested in our study (decades 2 – 10 | decade 3 is the reference: 20 – 30 years) and found that indeed, for both ProLong and RedLong genes, the mean expression incrementally reduces and increases, respectively, as we go from younger age groups towards the older age groups (Figure 4B). This difference across decades and types of genes is statistically significant (F = 32.405 | p-value = 8.900e^−41^). In addition to these trends, the expression of different genes from both the ProLong and RedLong groups also correlates with each other, as seen in the heatmap of the genes’ correlation (Figure 4C). All this data suggests that both ProLong and RedLong genes likely work together in a linked-regulatory fashion to orchestrate the threads of intrinsic skeletal muscle aging. This further corroborates that the ageprint is likely a set of co-expressed (and likely co-regulated) genes that drive baseline aging in a tissue.

**Figure 4:**
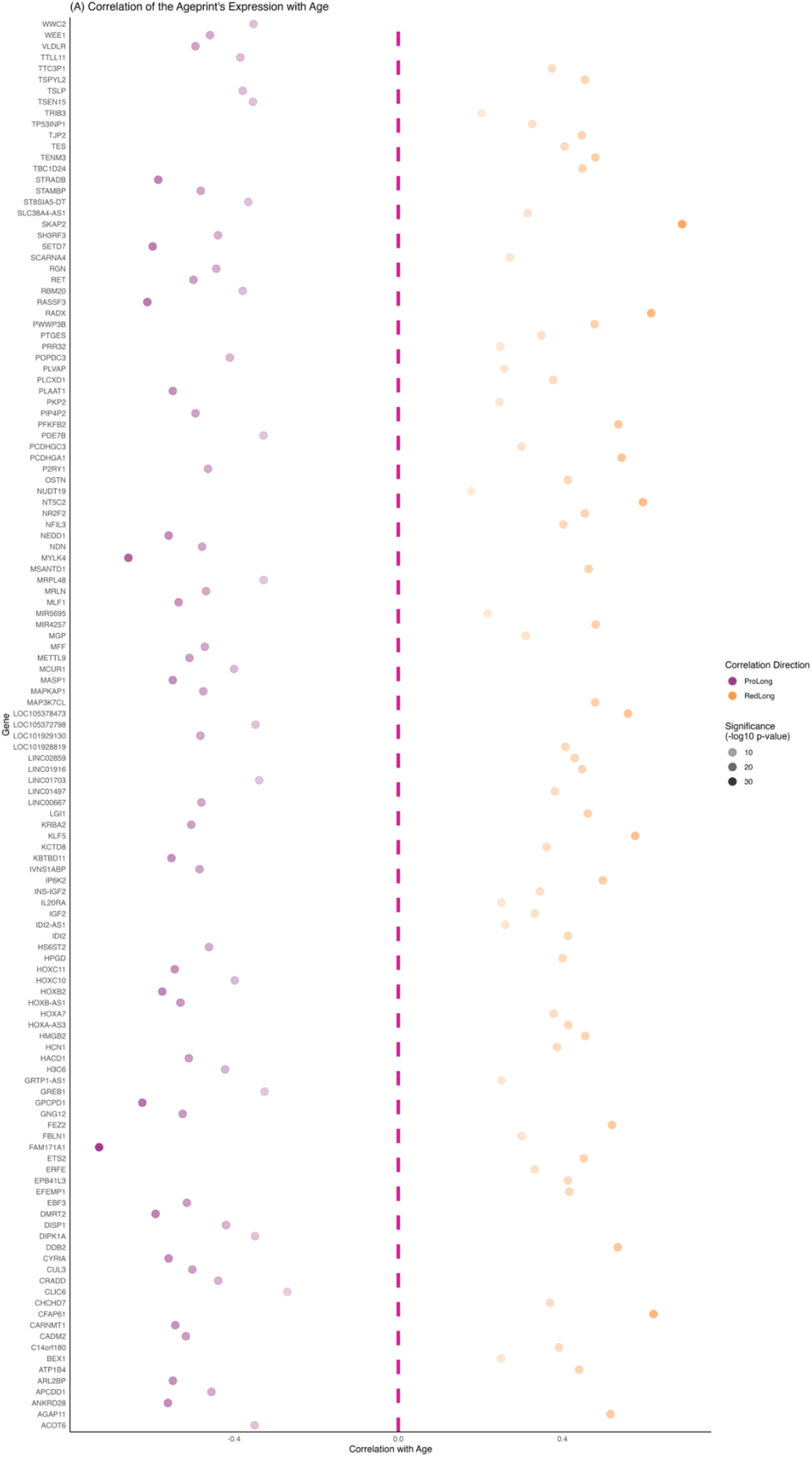
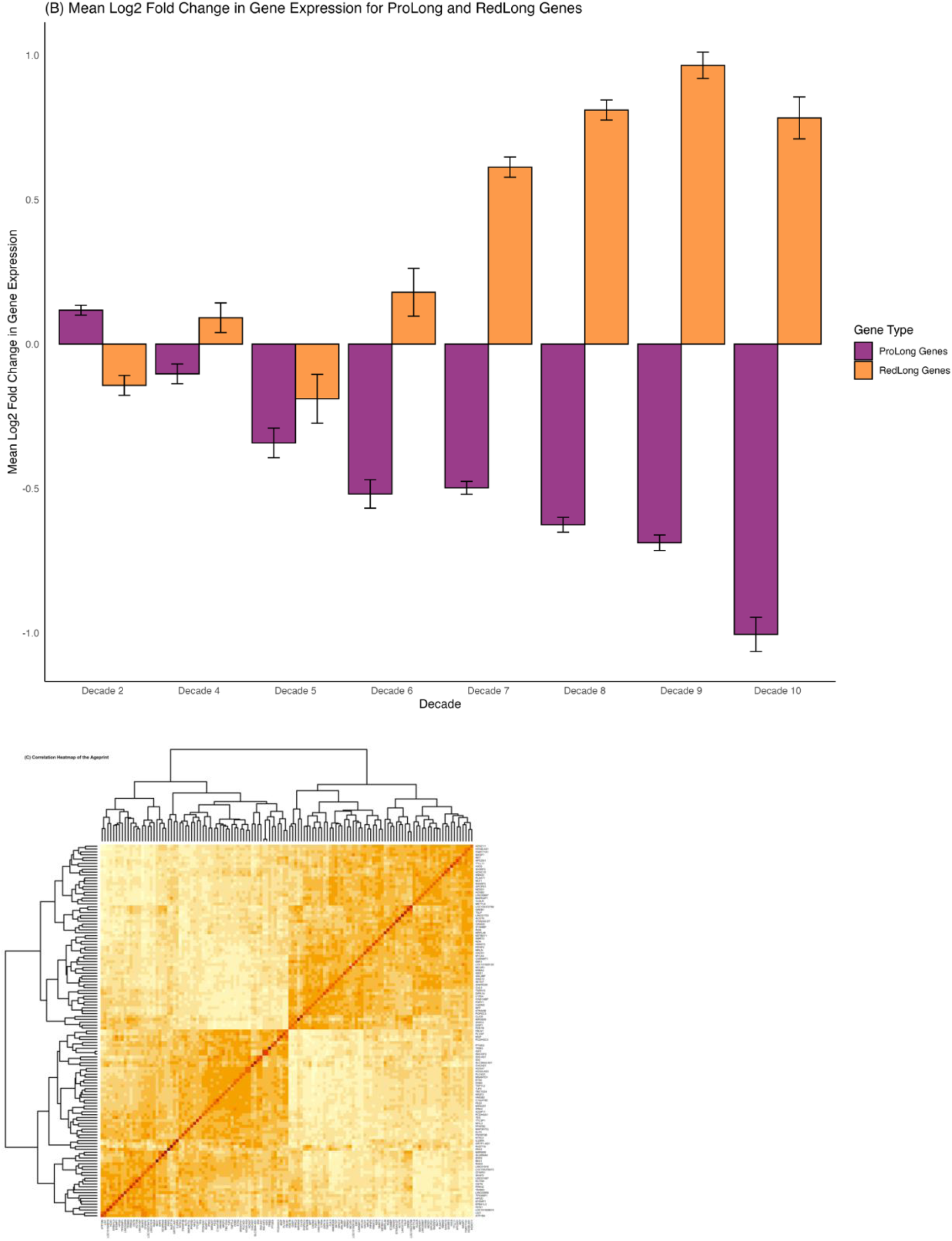
Correlation of the individual genes that constitute ageprint with chronological age. The genes in orange are RedLong genes, while the ones in purple are ProLong genes. A) shows the entire ageprint and the correlations of individual genes with chronological age. Surprisingly, all the genes negatively correlated with age are ProLong genes, while all the genes positively correlated with age are RedLong genes. B) shows the mean log fold changes in expression of the ProLong and RedLong genes across the different decades. Decade 3 (20 – 30 years) was chosen as the reference because muscle mass decreases once people are past their *prime*. C) is a heatmap of all the genes and their correlations. It shows that subsets of multiple genes’ expression correlate with each other, which further suggests that the genes are likely co-expressed and, potentially, even coregulated.

### Characterizing the ageprint genes

To understand the possible aging mechanisms of the ageprint genes, we looked at the molecular functions of the proteins they produce. We found that the functions of the genes involved a wide variety of cellular mechanisms, including epigenetic modifications, transcription regulation, DNA repair, and development. Several genes were also found to code for growth factor receptors, kinase domain-containing proteins, and zinc-finger proteins. An overview of some interesting genes and their functions is provided in Table 2. The presence of genes associated with growth factor receptors, insulin-like growth factors, kinases, PIP phosphatases, and NAD-binding proteins suggests that the ageprint of skeletal muscles largely influences the nutrient-sensing mechanisms of aging (Micó et al., 2017; J. Zhang et al., 2023). Moreover, the presence of HOX genes further corroborates our theory that the ageprint is a developmentally-linked genetic signature that regulates aging in a tissue. Our findings are further supported by evidence from the literature, which shows that HOX genes are associated with aging (D. Ko et al., 2024), including aging in skeletal muscles (Turner et al., 2020). Functional enrichment analyses did not yield tangible results since the number of genes was too small (128 genes).

**Table 2:** Ageprint genes grouped based on gene functions.

| Category | Genes | Notes |
| --- | --- | --- |
| **Methyltransferases** | METTL9, SETD7, CARNMT1 | Regulate the addition of methyl groups. |
| **Acetyltransferases** | ACOT6 | Regulates protein acetylation. |
| **Histones** | H3C6 | Chromatin protein. |
| **Transcription Factors** | ETS2, EBF3, DMRT2, BEX1, NFIL3 | Transcription regulation. |
| **DNA Repair** | PWWP3B, DDB2 | Regulating genome stability. |
| **Homeobox Genes** | HOXA7, HOXC11, HOXC10, HOXB2, HOXA-AS3, HOXB-AS1 | Linked to growth and development. |
| **Growth Factor Receptors** | EFEMP1, VLDLR, FBLN1, GREB1 | Likely control growth and nutrient sensing pathways. |
| **Insulin/IGF Genes** | INS-IGF2, IGF2 | Key regulators of metabolic processes. |
| **Phospholipases** | PLCXD1, PIP4P2 | May regulate PIP2 levels, affecting cell signaling. |
| **mTOR Complex Members** | MAPKAP1 | Could be involved in nutrient sensing and aging pathways. |
| **Kinase Domain Proteins** | TJP2, TRIB3, WEE1, RET, IP6K2, STRADB, MYLK4, DIPK1A, PFKFB2 | Regulators of phosphorylation and signaling pathways. |
| **Zinc Finger Proteins** | KLF5, TES, SH3RF3, KRBA2, NR2F2, RBM20, IVNS1ABP | Involved in DNA binding and transcriptional regulation. |
| **Immunoglobulin Domains** | EBF3, GPCPD1, CADM2, IL20RA | Likely linked to cell-to-cell signaling. |
| **Senescence Marker** | RGN | Known marker of cellular senescence. |
| **NAD-Binding Proteins** | CFAP61, HPGD | Can be involved in metabolism and aging. |
| **Non-Protein Coding Genes** | MIR3659, LINC01703, LINC00667, MIR4257, LINC02859, LINC01497, LINC01916, MIR5695 | Non-coding RNAs. |

### Novel targets are related to nucleotide and fatty acid metabolism

We performed GO-term enrichment to understand the biological processes associated with the 26 novel targets identified by SkeletAge (Figure 5). Most GO-terms can be categorized into two distinct processes: nucleotide and fatty acid metabolism. This suggests that these novel targets, constituents of skeletal muscles’ ageprint, are linked to nucleotide and fat metabolism. Out of the 26 novel targets, 16 are predicted to increase longevity (ProLong genes), while the remaining 10 are predicted to reduce longevity (RedLong genes). Most GO-terms associated with the 16 novel ProLong genes seem to be involved in nucleotide metabolism. This suggests that the genes that regulate nucleotide metabolism in skeletal muscles show downregulation with age, given how all the ProLong genes were negatively correlated with age and showed a consistent, incremental decline in their expression. In fact, sarcopenia, the characteristic decline in muscle mass, volume, and strength, is a hallmark of skeletal muscle aging (Yaku et al., 2018) that may be linked to changes in cellular adenine metabolism (Frederick et al., 2016). This suggests that addressing some of the novel targets discovered by SkeletAge might help overcome declining NAD levels in aging muscle tissue (Chubanava & Treebak, 2023). On the other hand, the 10 novel targets predicted to be involved in reducing longevity (RedLong genes), are linked to GO-terms that suggest an involvement in fatty acid metabolism. Since all RedLong genes were positively correlated with age, it seems that genes that regulate fatty acid metabolism in muscles get upregulated over time. Increased fat accumulation in the muscles (myosteatosis) is a pathology that is distinct from sarcopenia (Ahn et al., 2021; Miljkovic & Zmuda, 2010; L. Wang et al., 2024). Myosteatosis is recognized as a common occurrence in aging, which leads to reduced muscular strength (Biltz et al., 2020) and altered muscle architecture (Jiang et al., 2019) that ultimately adds to age-related musculoskeletal phenotypes. Therefore, targeting these novel RedLong genes might help overcome some of the myosteatosis-associated phenotypes in aging muscles.

**Figure 5:**
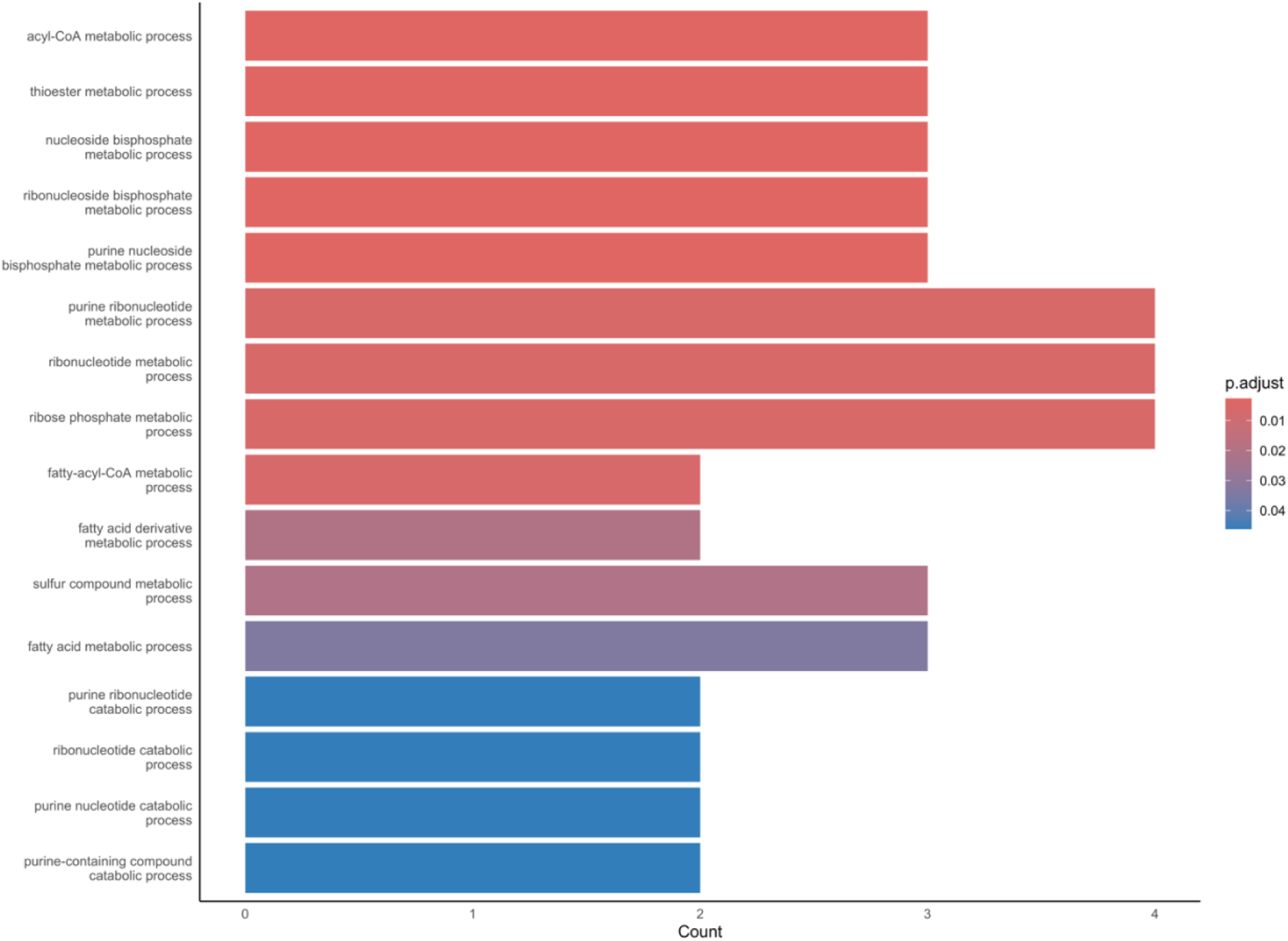
GO-terms enriched for the 26 novel targets identified using SkeletAge. Overall, the terms can be categorized into two major processes: nucleotide and fatty acid metabolism. Both of these processes contribute to age-associated muscle loss, which further demonstrates that identifying the ageprint can shed light on the mechanisms of baseline, healthy aging across tissues.

### Finding druggable targets to discover pharmacological interventions for aging

To see if any of the genes identified by SkeletAge were druggable targets, we ran the ageprint on DGIdb (Cannon et al., 2024; Griffith et al., 2013) and filtered the hits for all the genes with an interaction score of > 1.0 (Figure 6). By doing so, we found that 23 of the ageprint’s constituent genes were part of the druggable genome (Hopkins & Groom, 2002; Russ & Lampel, 2005), and 9 of those genes had approved drugs listed against their protein products (Table 3); one of these 9 genes (*PDE7B*) was among the 26 novel targets discovered by SkeletAge. Interestingly, out of the 9 genes with approved drugs, 3 genes (*PDE7B*, *NT5C2*, *TRIB3*) were associated with drugs that have previously been investigated in aging. *PDE7B* was associated with dyphylline, which was recently shown to improve fertility in naturally-aged mice by increasing primordial follicle activation – the results were also replicated in human ovarian cells (W. Zhang et al., 2024). Didanosine, a reverse transcriptase inhibitor linked to *NT5C2*, was shown to extend the lifespan of *C. elegans* by almost 28% (Brochard et al., 2023; McIntyre et al., 2023). Indapamide and perindopril were both linked to *TRIB3*. Indapamide, which can cross the blood-brain barrier, demonstrated neuroprotective and antioxidative effects against age-associated lipid peroxidation and loss of myelin and axons in middle-aged mice (Michaels et al., 2020). Perindopril was directly investigated in older people who had problems related to physical mobility in a clinical trial (International Standard Randomised Controlled Trial Register: ISRCTN67679521) and was found to improve 6-minute walking distance, lower blood pressure, and lead to an overall better self-reported quality of life. The effect of perindopril on the improvement in physical capacity was comparable to 6 months of exercise for the elderly (Sumukadas et al., 2007). This suggests that the ageprint predicted by SkeletAge very likely contains genes that influence and regulate the mechanisms of baseline aging in skeletal muscles. In the future, we will hopefully see more potential drugs listed in Table 3 being studied in the context of skeletal muscle aging. These findings underscore the importance of identifying a tissue’s ageprint for longevity drug discovery.

**Figure 6:**
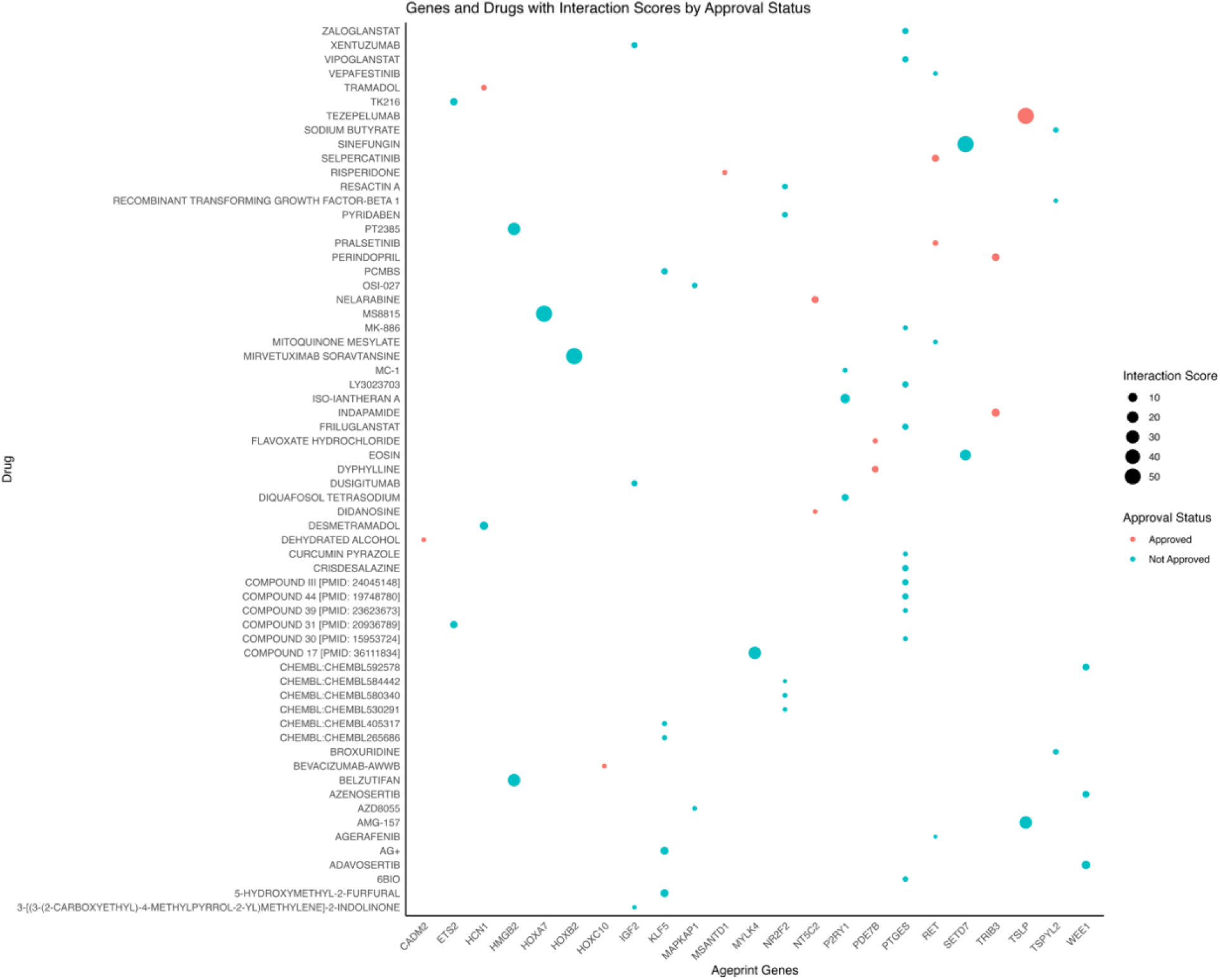
A scatterplot of all the ageprint genes that are part of the druggable genome and their associated drugs. The size of the points reflects the interaction score. PTGES has the most approved drugs associated with it, followed by *NR2F2* and *KLF5*.

**Table 3:** The 9 genes that are a part of the ageprint and have approved drugs listed against their protein products. 4 drugs on this list have already been investigated in the context of aging and found to have promising results.

| Gene | Drug |
| --- | --- |
| CADM2 | Dehydrated alcohol |
| HCN1 | Tramadol |
| HOXC10 | Bevacizumab-awwb |
| MSANTD1 | Risperidone |
| NT5C2 | Didanosine; nelarabine |
| PDE7B | Flavoxate hydrochloride; dyphylline |
| RET | Selpercatinib; pralsetinib |
| TRIB3 | Indapamide; perindopril |
| TSLP | Tezepelumab |

### Skeletome: a skeletal muscle database to accelerate aging research

The first step of building SkeletAge was identifying the right datasets for the training and validation set. Given the stringent requirements for our inclusion criteria, we had to gather a lot of publicly available RNA-seq data from GEO. One of the main challenges was acquiring accurate age labels for different samples because most groups had reported average ages for different groups in their papers. Thus, we reached out to several authors and corresponding principal investigators on the papers to get information about the individual samples’ ages and then put it all together manually. Over time, we curated a large database of 534 samples in total, which pulls together data from over 11 different studies, including a couple of clinical trials – we call this database *Skeletome*. The samples in the Skeletome range from 19 – 90.7 years, with an average age of 52 years and a standard deviation of ±21.3 years (Figure 7), thereby covering a large enough proportion of ages to measure genetic effects across the individuals ranging from young adults to almost centenarians. Additionally, Skeletome comprises data from studies that investigated the pre- and post-intervention effects of different “anti-aging” supplements and lifestyle patterns such as nicotinamide mononucleotide (NMN), high-intensity workout, and even sedentary lifestyle. Moreover, it also has data on various health conditions that may accelerate biological aging, such as obesity (Foster et al., 2023; Lin et al., 2021) and type-2 diabetes (Fraszczyk et al., 2022). All the data in Skeletome comes from RNA-seq performed on biopsies of vastus lateralis tissue, thus ensuring homogeneity and reducing systematic errors and confounders arising from differences in skeletal muscle types. In addition to the metadata, Skeletome contains the raw counts for each sample, aligned to the GRCh38, thereby making it simpler (and faster) to study gene expression in skeletal muscle tissue across different ages. In compiling Skeletome, we hope this database would further accelerate research in skeletal muscle genomics by removing a key bottleneck: the time it takes to reinvent the wheel by individually reaching out to every team and getting age labels. You can access the Skeletome data through SkeletAge’s GitHub repository: https://github.com/mali8308/SkeletAge/.

**Figure 7:**
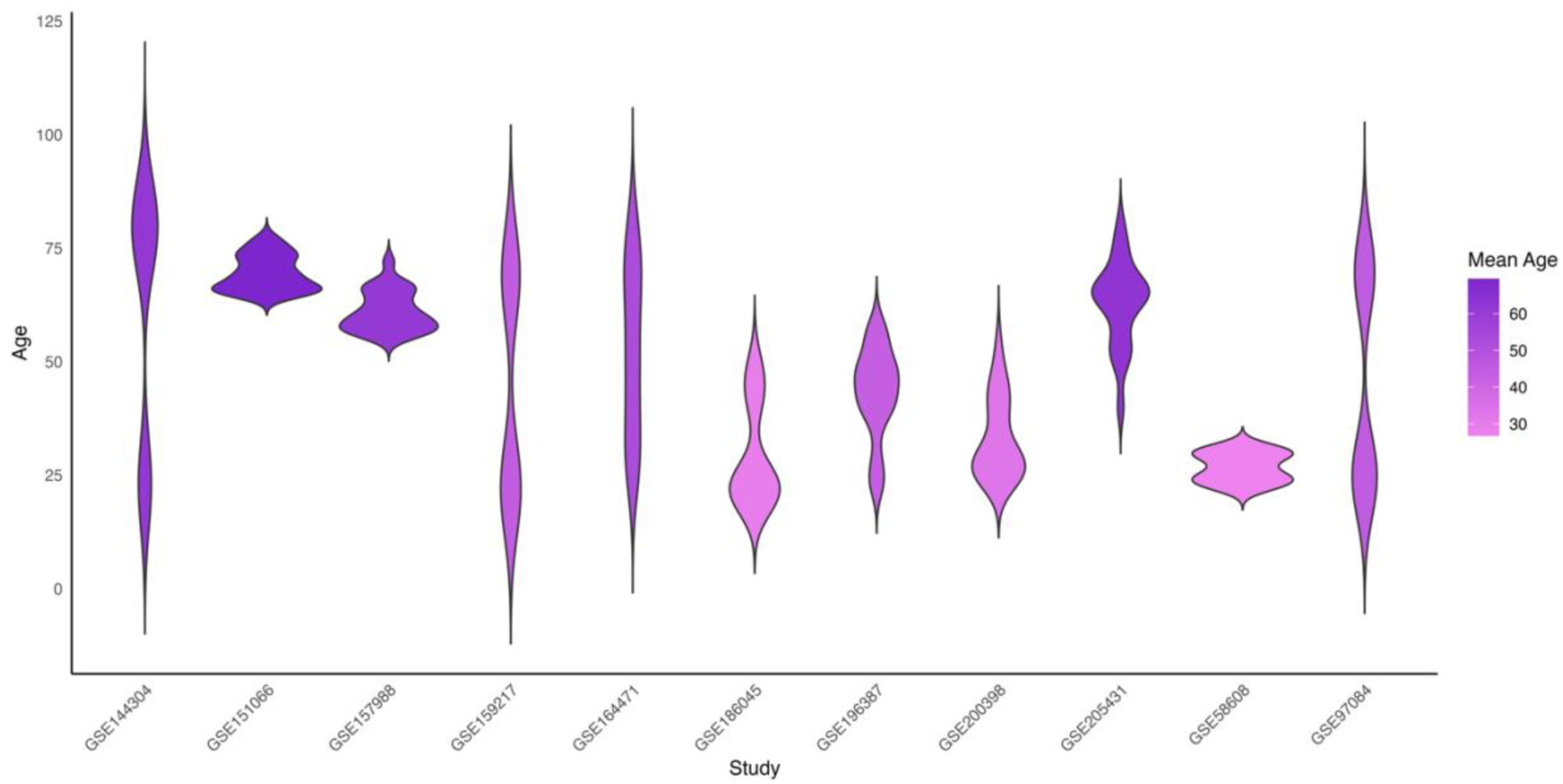
A violin plot showing the chronological age distribution across Skeletome. Different datasets have different age distributions. Each dataset’s mean age is reflected in the intensity of the color. The curve’s width is proportional to the number of samples for that particular age range in a dataset.

## Discussion

Recent paradigm shifts in geroscience research have created opportunities to address aging as a disease and investigate it like one. While identifying disease-causing genes is usually one of the first steps for tackling non-infectious diseases, genes that cause aging have remained rather elusive in humans. Thus, in this paper, we describe a method to identify the *ageprint* of a tissue – the tissue-specific set of genes that regulate developmentally-linked baseline “healthy” aging, i.e., aging that occurs when there is no disease or aging interventions at play – using a transcriptomics-based aging clock called *SkeletAge*. Using SkeletAge we narrowed down the skeletal muscle ageprint to 128 genes; 26 of these genes have never been studied in the context of aging or aging-associated phenotypes. Additionally, 23 ageprint genes were found to be a part of the druggable genome. Hence, we demonstrate how identifying the ageprint of a tissue can enhance our understanding of the mechanisms that drive aging and accelerate the development of pharmacological interventions against aging.

In skeletal muscles, SkeletAge performs better than a previously built tissue-specific age-estimator, RNAAgeCalc (Ren & Kuan, 2020). We believe that this difference in performance can be accounted for by the discrepancy in the training sets of SkeletAge and RNAAgeCalc. SkeletAge was built using data from healthy people who were alive at the time of sample collection, whereas RNAAgeCalc relies on post-mortem data from GTEx. Thus, we also highlight a new criterion for choosing the training set when building a transcriptomics-based age-estimator to identify the ageprint of a tissue.

Based on GO-term enrichment, the 26 novel targets identified by SkeletAge seem to be involved in nucleotide metabolism and myosteatosis. There is a marked decrease in NAD levels with age (Chubanava & Treebak, 2023). A study found that knocking out Nampt – an enzyme that helps produce NAD and, consequently, ATP – leads to progressive muscle weakness and a decline in physical fitness as mice age. Moreover, they also found that the weakened phenotype can be rescued through NR supplementation, a compound that bypasses the need for the Nampt enzyme. Additionally, the team overexpressed Nampt in the skeletal muscles and found that it preserves some endurance in older mice (Frederick et al., 2016). This suggests that NAD and its precursors might help in overcoming some of the age-associated skeletal muscle dysregulation, as they have in yeast (Odoh et al., 2022; Wilson et al., 2023). However, as in the clinical trial NCT03151239 (Yoshino et al., 2021), we found that NAD did not significantly lower the biological age of skeletal muscles. This suggests that while the genes linked with NAD metabolism contribute to aging in skeletal muscles, their (dys)function likely cannot be rescued by NMN supplementation alone. Myosteatosis, the increased intramuscular accumulation of fats, is another phenotype associated with skeletal muscle aging (Imamura et al., 1983), that is not linked to sarcopenia (Ahn et al., 2021; Miljkovic & Zmuda, 2010; L. Wang et al., 2024). Perhaps studying the novel targets identified by SkeletAge might help elucidate how to regulate or reduce myosteatotic changes in muscles as we age. The ability of SkeletAge to identify a group of genes that not only precisely predict and (as a set) strongly correlate with aging but also have functions that reflect known phenotypes associated with skeletal muscle aging suggests that it is correctly identifying the ageprint. This is further corroborated by the fact that using the ageprint, SkeletAge can accurately assess the effect of longevity therapeutics on aging.

Some of the genes involved in the ageprint were also a part of the druggable genome (Hopkins & Groom, 2002; Russ & Lampel, 2005). 9 of these genes were associated with approved drugs, and out of those 4 drugs (dyphylline, didanosine, indapamide, perindopril) had already been tested in the context of aging. Dyphylline improved fertility in aged mice (*in vivo*) and human oocytes (*in vitro*) by activating primordial follicles by inhibiting phosphodiesterases that break down cAMP (W. Zhang et al., 2024). This leads to the activation of the PI3K/Akt pathways, which results in the translocation of FOXO3 to the cytoplasm and the activation of follicles (Ting & Zelinski, 2017). Thus, dyphylline might help in rescuing fertility in older women who face ovarian insufficiency, which can lead to premature menopause (Cox & Liu, 2014). Didanosine was recently shown to be able to extend the lifespan of *C. elegans* (Brochard et al., 2023; McIntyre et al., 2023). Didanosine is a reverse-transcriptase inhibitor that likely causes mitochondrial toxicity, which leads to mitohormesis – a process whereby mitochondrial damage leads to the upregulation of cellular defense mechanisms, including the overexpression of ATF-4, a transcription factor that helps cells respond to stress by inhibiting translation (Costa-Mattioli & Walter, 2020) and has been shown to increase lifespan in worms (Statzer et al., 2022). This suggests that the use of didanosine (or other mild mitotoxic drugs) might lead to preferential expression of stress-response genes (J. Wang et al., 2010), and the increased translation of protective proteins (Rogers et al., 2011) through the expression of ATF-4, which might culminate in a net protective response that may extend lifespan (Brochard et al., 2023). Indapamide, a diuretic used to treat hypertension (Fiddes et al., 1997; Syed, 2022), was found to have neuroprotective and antioxidative effects against age-induced oxidative damage and myelin loss in mice (Michaels et al., 2020). With age, reactive oxygen species (ROS) production increases in microglia (O’Neil et al., 2018), likely through the increased expression of NADPH oxidase, which can lead to tissue damage. Indapamide, which has antioxidant properties (Tamura et al., 1990) and can cross the blood-brain barrier, has been shown to reduce axon loss and demyelination in middle-aged mice, which is associated with reduced oxidative stress (Michaels et al., 2020). This suggests that indapamide and other similar antioxidants may offer neuroprotection against myelin and axon loss as we age. Perindopril was investigated in humans in a clinical trial (ISRCTN67679521) and was found to improve 6-minute walking distance in elderly people by about 30 meters – this is comparable to over 6 months of exercise. Perindopril is an angiotensin-converting-enzyme (ACE) inhibitor, a class of drugs used to treat COVID-19 with mixed results (Hippisley-Cox et al., 2020; Najafi et al., 2022; Writing Committee for the REMAP-CAP Investigators, 2023). ACE inhibitors have previously been linked to a slower decline in muscular strength in elderly women (Onder et al., 2002). Moreover, using perindopril also helped improve the 6-minute walking distance in elderly people who had experienced heart failure (Hutcheon et al., 2002). This suggests that perindopril could likely improve vascular function in the elderly and help slow aging. Thus, these findings suggest that the remaining approved drugs that were linked to the ageprint (dehydrated alcohol, selpercatinib, pralsetinib, tramadol, risperidone, flavoxate hydrochloride, bevacizumab-awwb, nelarabine, tezepelumab), might also have geroprotective mechanisms that could help alleviate some of the phenotypes associated with aging.

One of the main limitations of using the ageprint to study aging is its tissue-specificity, which means it cannot be generalized as a universal signature of aging. However, recent evidence has shown that different tissues age at different rates (Oh et al., 2023; Slieker et al., 2018; Tian et al., 2023; Tuttle et al., 2020), so it might make more sense to target each tissue’s aging individually. While this approach is not perfect, we believe that by finding the ageprint for every tissue, we can find the *universal ageprint* for humans at the intersection of all the ageprints. That universal signature of healthy aging might help us develop drugs that simultaneously target aging at the molecular level in multiple tissues. Thus, in the future, we hope to dissect this universal ageprint further and get closer to understanding the basic mechanisms of aging.

Finally, our findings demonstrate that using tissue-specific age predictors to identify the ageprint is an efficient way of discovering tissue-specific novel targets for aging research, which can inform both the mechanistic and the drug-discovery side of aging research.

## Methods

### Training set inclusion criteria

To capture the ageprint of skeletal muscles, we wanted to train our elastic net model on vastus lateralis samples taken from healthy individuals over a broad age range. We hypothesized that genes that can explain healthy aging at younger and older ages would constitute the true ageprint of a tissue. We decided on 236 total samples from GSE164471, GSE97084, and GSE144304. The data in GSE164471 came from the participants of the GESTALT study (Tumasian et al., 2021). The study only included participants who met the IDEAL (Insight into the Determination of Exceptional Aging and Longevity) standard developed by the National Institute on Aging (Schrack et al., 2014). The participants had no chronic illnesses (except for hypertension and cancer that has not relapsed for the last 10 years), that they took medication for (except for antihypertension medication), no prior history of professional training, had a BMI < 30 kg/m^2^, and were not diagnosed with any physical impairment, major disease, or cognitive damage. For the data taken from GSE97084, participants were only included if they did not work out regularly (more than 20 min, twice a week), had no cardiovascular problems, or metabolic disorders such as hypo- or hyperthyroidism, type-2 diabetes (fasting blood glucose > 110 mg/dL), and their BMI ≤ 32 kg/m^2^. Participants were excluded if they had metal implants, suffered from renal diseases, were pregnant, or had a history of blood clotting problems. Participants were also excluded if they used any of the following medications: corticosteroids, tricyclic antidepressants, sulfonylureas, barbiturates, peroxisome proliferator-activated receptor γ agonists, anticoagulants, β blockers, opiates, insulin, and insulin sensitizers (Robinson et al., 2017). The data from GSE144304 comes from the FITAAL study (Dutch Trial Registry identifier: NTR6124). Participants were not included in the trial if they had COPD, cancer, dementia, cardiac failure, neuromuscular disorder, anemia, or were contraindicated to receive a muscle biopsy. They were also excluded if they were currently part of another study or had recently undergone an intensive medical treatment or procedure, such as surgery. Other physiological parameters that could get participants excluded were a BMI <20 kg/m^2^ or >25 kg/m^2^, having type I or II diabetes mellitus, regular workout routine (more than 4 times a week), or pregnancy and/or nursing states (de Jong et al., 2023). The overarching trend for sample selection across the three different datasets was healthy people with a BMI >20 kg/m^2^ or >32 kg/m^2^ who were not professional athletes and did not work out regularly, were free of any major illnesses or chronic conditions that would require medication and/or treatments which can – in-itself or together with the condition – confound the gene expression patterns, and were free of physical or cognitive impairments.

### Creating the Skeletome Database

To study gene expression changes in skeletal muscle, we first created a large data set of 534 vastus lateralis samples from 11 different studies on NCBI Gene Expression Omnibus (GEO): GSE164471 (Tumasian et al., 2021), GSE97084 (Robinson et al., 2017), GSE144304 (de Jong et al., 2023), GSE196387 (Whytock et al., 2023), GSE205431 (Drummer et al., 2022), GSE157988 (Yoshino et al., 2021), GSE151066 (Rubenstein et al., 2022), GSE58608 (Lindholm et al., 2014), GSE159217 (Lagerwaard et al., 2021), GSE200398 (Semenova et al., 2022), GSE186045 (McFarland et al., 2022) – for all of these studies, we extracted their metadata and the raw counts directly from NCBI GEO. The reads are aligned using HISAT2 (Kim et al., 2019) to GCA_000001405.15. Runs with an alignment rate of over 50% are used to generate the counts matrix using Subread featureCounts, and the *Homo sapiens* Annotation Release 109.20190905 is used for gene annotation. Wherever necessary, we reached out to the groups that had published these results and got individual age labels for samples to map accurate ages for the penalized regression model training and testing. The raw counts and the metadata (including the GEO sample identifier, accurate age label, and condition) are present in a database called Skeletome. We hope that Skeletome on its own would help accelerate aging research in skeletal muscle by providing a substantial dataset for future transcriptomics-based clocks, aging therapy development, and more. Skeletome can be accessed through our GitHub repository: https://github.com/mali8308/SkeletAge/.

### DESeq2 for normalization and expression tracking

To process raw counts, we used DESeq2 (Love et al., 2014). We divided our samples into bins of 10 years, which started from decade 1 (0-10 years) and went all the way up to decade 10 (90-100 years). This made it easier to create a factor with 11 different levels of comparison instead of 116 different levels that we would have had to use if we had chosen individual sample ages as the factors. We chose the third decade (20-30 years) as the reference for DESeq2 because muscle mass tends to start declining as people enter ages beyond 30 years (Lexell et al., 1988) and may reduce by over 40% as they grow past 80 years of age. Since we wanted to capture healthy aging, we used the “prime” age as the reference to compare gene expression against. Moreover, biological age measurements are often non-linear during early growth and development (Horvath & Raj, 2018), which means that choosing a younger reference level (people who were younger than decade 3) could have confounded the skeletal muscle ageprint that we were trying to detect.

### Glmnet for training SkeletAge and feature selection

SkeletAge is based on an elastic net regression model (Zou & Hastie, 2005), which helps select important aging genes that form a tissue’s ageprint. We used the glmnet package (4.1-8) in R to generate an elastic net regression model using leave-one-out cross-validation for hyperparameter tuning (λ = 19.15027 | α = 0.1). We trained SkeletAge on the samples in the following studies: GSE164471, GSE97084, and GSE144304. Using a penalized regression model automatically reduced the coefficients for all the genes that were not important down to 0. Thus, to extract the ageprint from SkeletAge, we selected all the β ≠ 0, except the intercept (β_0_). Using this strategy gave us 128 genes that seem to be central to skeletal muscle aging: the *ageprint*.

### Validating SkeletAge’s performance

To ensure that SkeletAge was truly capturing the ageprint of skeletal muscles, we created a validation set using data from 8 different studies (GSE196387, GSE205431, GSE157988, GSE15106, GSE58608, GSE159217, GSE200398, and GSE186045). The samples in these studies range from extremely healthy (such as athletes) to people who are overweight or obese, have type-2 diabetes, obesity, and more. We wanted to create a diverse set so that their condition would influence their gene expression. This would allow us to test whether the ageprint gets affected by health status by determining whether SkeletAge can still precisely map the correct ages for the samples. Theoretically, a robust ageprint would allow SkeletAge to predict precise ages despite the altered state. Our results demonstrate that even for the diverse dataset, SkeletAge could use the skeletal muscle ageprint to predict ages that were not statistically different from the samples’ chronological ages. Being able to do that using the 128 genes we have identified suggests that these genes are a universal signature that remains unchanged regardless of health status and, hence, play a pivotal role in driving healthy skeletal muscle aging.

### Comparison against RNAAgeCalc and statistical analysis

To benchmark SkeletAge’s performance, we tested it against an already-published RNA-seq-based clock that allows you to specify the tissue for which you would like to predict age: RNAAgeCalc (Ren & Kuan, 2020). We used the RNAAgeCalc package in R and used the predict_age function. We passed it the raw counts of both our training and validation set, with the “tissue” and “exprtype” arguments set to “muscle” and “counts,” respectively. For statistical tests, we ran Wilcoxon signed rank tests in R at the alpha of 0.05. We measured the difference between the chronological ages of the sample and the ages predicted by SkeletAge, the chronological ages of the sample and the ages predicted by RNAAgeCalc, and the difference between the ages predicted by SkeletAge and RNAAgeCalc. We also computed the Pearson correlation coefficient between the chronological ages and the ages predicted by SkeletAge and the chronological ages and the ages predicted by RNAAgeCalc. We also calculated the coefficients of determination (R^2^) for both SkeletAge and RNAAgeCalc on both the training set and the validation set.

### Enrichment analysis, finding druggable targets, and generating plots

To retrieve information about genes, their descriptions, names, InterPro accession numbers, and InterPro descriptions, we used biomaRt (3.19) (Durinck et al., 2005). We used the clusterProfiler (Yu et al., 2012) package in R for enrichment analyses, including GO-term enrichment, reactome enrichment, disease enrichment, and KEGG enrichment. We also used STRINGdb (Szklarczyk et al., 2022) version 12 for protein-protein interaction analysis and networks. Due to the relatively small number of genes, we did not see any enrichment for reactome and KEGG or significant protein-protein interactions with STRINGdb. We used the DGIdb (Cannon et al., 2024; Griffith et al., 2013) to identify the constituent genes of the ageprint, which were also druggable targets. Lastly, we used the ggplot2 package (3.5.1) in R to create all the plots used throughout the paper.

## Abbreviations

RNA-seq: RNA-sequencing
CI: Confidence intervals
MAE: Mean absolute error

## Author Contributions

M.A. and M.K. developed the SkeletAge estimator. M.A. conceived the study and performed the statistical analysis. M.A., F.L., and M.K. wrote the manuscript.

## Acknowledgement

We would like to thank the following people for sharing accurate age labels for the samples which ultimately helped us in building and validating SkeletAge:

- Center for Human Nutrition, Washington University School of Medicine: Samuel Klein, Gordon Smith.
- Division of Cardiovascular Medicine, Stanford University School of Medicine: Maléne Lindholm.
- Department of Physiology and Pharmacology, Karolinska Institutet: Carl J. Sundberg.
- Mayo Clinic: Surendra Dasari, K. Sreekumaran Nair.
- College of Health, Oregon State University: Matthew Robinson.
- Wageningen University: Evert van Schothorst, Jaap Keijer.
- Translational Research Institute, AdventHealth: Katie Whytock, Lauren Sparks, and Paul Coen.

## Supplementary Material

**Table S1: Details about the datasets that were used to create and validate SkeletAge**. The table contains the NCBI GEO accession number, the general phenotype of the individuals whose data was collected, the number of samples taken from each dataset, and whether that dataset was used to train or test SkeletAge. All of these samples have had bulk RNA-seq performed on vastus lateralis biopsies.

**Table S2: The genes that make up the ageprint of skeletal muscles**. Out of these 102 genes have been already studied in the context of aging. The remaining 26 are present in table 1.

**Table S3: Ageprint genes and their association with aging in the literature**. This table presents sources where individual or multiple ageprint genes are studied in the context of aging or age-associated phenotypes.

